# Distinct cell types along thick ascending limb express pathways for monovalent and divalent cation transport

**DOI:** 10.1101/2025.01.16.633282

**Authors:** Hasan Demirci, Jessica Bahena-Lopez, Alina Smorodchenko, Xiao-Tong Su, Jonathan Nelson, Chao-Ling Yang, Joshua Curry, Xin-Peng Duan, Wen-Hui Wang, Yuliya Sharkovska, Ruisheng Liu, Duygu Elif Yilmaz, Catarina Quintanova, Katie Emberly, Ben Emery, Nina Himmerkus, Markus Bleich, David H. Ellison, Sebastian Bachmann

**Author notes:** Correspondence: Prof. Dr. Sebastian Bachmann Charité-Universitätsmedizin Berlin Institut für Zell-und Neuroanatomie Philippstr. 12, 10115 Berlin, David H. Ellison, MD, Division of Nephrology & Hypertension Oregon Health & Science University Portland, OR 97239, Oregon. H.D. and J.B.L. contributed equally to this work. Equal corresponding author (David H. Ellison, Sebastian Bachmann). Competing interests: The authors declare no competing interest.

## Abstract

Kidney thick ascending limb cells reabsorb sodium, potassium, calcium, and magnesium and contribute to urinary concentration. These cells are typically viewed as of a single type that recycles potassium across the apical membrane and generates a lumen-positive transepithelial voltage driving calcium and magnesium reabsorption, although variability in potassium channel expression has been reported. Additionally, recent transcriptomic analyses suggest that different cell types exist along this segment, but classifications have varied and have not led to a new consensus model. We used immunolocalization, electrophysiology and enriched single nucleus RNA-Seq to identify thick ascending limb cell types in rat, mouse and human. We identified three major TAL cell types defined by expression of potassium channels and claudins. One has apical potassium channels, low basolateral potassium conductance, and is bordered by a sodium-permeable claudin. A second lacks apical potassium channels, has high basolateral potassium conductance and is bordered by calcium- and magnesium-permeable claudins. A third type also lacks apical potassium channels and has a high basolateral potassium conductance, but these cells are ringed by sodium-permeable claudins. The recognition of diverse cell types resolves longstanding questions about how solute transport can be modulated selectively and how disruption of these cells leads to human disease.

## Introduction

The kidney thick ascending limb (TAL) reabsorbs several filtered solutes and also contributes importantly to urinary concentration and dilution. As physiological challenges to extracellular fluid (ECF) volume, divalent ion metabolism, or water balance may occur independently, the TAL must be able to adjust solute transport selectively in response to physiological need. Yet, TAL cells are generally viewed as expressing a common set of key transport proteins. These include the sodium, potassium, chloride transporter (NKCC2) and a K^+^ channel (ROMK) at the apical membrane, and a chloride channel at the basolateral membrane. The combination of K^+^ recycling across the apical membrane and Cl^-^ exit across the basolateral membrane generates a transepithelial voltage oriented with the lumen positive relative to interstitium (1–3).

Ca^2+^ and Mg^2+^ reabsorption along the TAL take place via a paracellular pathway comprising claudins 16 and 19 (4), driven by the transepithelial voltage. Transport of these ions can be modulated separately from that of Na^+^. High Ca^2+^ intake, for example, decreases paracellular Ca^2+^ transport along the cortical TAL (cTAL) without altering transcellular NaCl transport (5), an effect that is likely mediated in part by the calcium sensing receptor (CaSR) (6). While activating mutations of the CaSR in humans sometimes lead to mild salt wasting (7), the bulk of evidence indicates that CaSR activation along the TAL primarily affects Ca^2+^ transport with minimal effects on transepithelial voltage or NaCl excretion (6, 8). Along the same lines, parathyroid hormone (PTH) appears capable of stimulating Ca^2+^ reabsorption with little or no effect on transepithelial voltage or NKCC2 (9).

One way that differential regulation may occur is because the TAL comprises different cell types. Early studies in amphiuma and hamster posited the existence of two cell types, one with high basolateral K^+^ conductance (HBC) and another with low basolateral conductance (LBC) (10, 11). It was later shown that some TAL cells don’t express ROMK at their apical membrane (12, 13), a finding incorporated by Dimke and Schnermann in a TAL model with two cell types based on K^+^ conductance (14). Yet, their model did not link K^+^-conductive cell types to cells that transport Ca^2+^ and Mg^2+^.

More recently, Milatz and colleagues (15) reported that different claudins, tight junction proteins that act as paracellular solute ‘gates’, are expressed in a mutually exclusive *mosaic pattern* along the TAL. As these claudins are selectively permeable for divalent (claudins 16 and 19) and monovalent cations (claudin 10), this suggested that there are discrete paracellular pathways for the different cations (4). Recently, transcriptomic analyses have also detected clusters of TAL cells, one predominantly expressing claudin 16 and the other predominantly expressing claudin 10 (16–18), seemingly aligning with the immunolocalization results. Yet the terms used to denote cells and the gene signatures involved (17–20) have not been integrated with important, but often neglected, earlier results concerning sodium and potassium transport pathways (10–13). Here, by combining immunocytochemical, transcriptomic and electrophysiological approaches across multiple species, we show the existence of two primary TAL cell types in the cortex and outer stripe of the outer medulla (OSOM), one that should transport Na^+^ efficiently and generate the transepithelial voltage while the other should be electroneutral, transport Ca^2+^ and Mg^2+^, and respond to regulators of Ca^2+^ metabolism. A third cell type, present only in the inner stripe of the outer medulla (ISOM), combines key features of the other two and likely participates in K^+^ reabsorption.

## Results

### Immunohistochemistry Reveals Mosaicism of Ion Channel Expression

We first confirmed that, in contrast to uniform expression of NKCC2 along the entire TAL, apical immunoreactive ROMK signal shows mosaic expression along both medullary (m)TAL and cTAL (**Figure 1A, Supplemental Figure 1A**). ROMK signal was concentrated in the apical aspect of cells categorized as *positive* with primary antibody concentration adjusted to optimal signal-to-noise ratio. In this standard condition, cells categorized *negative* were lacking luminal ROMK signal. At electron microscopic (EM) resolution, ROMK immunoreactive signal was strictly located at the apical membrane of ROMK-positive TAL cells by immunoperoxidase staining; the signal ended abruptly at the tight junctions coupling ROMK-positive to adjacent ROMK-negative cells (**Figure 1B and C**). cTAL segments contained 28 ± 2% ROMK-negative cells, medullary (m) TAL of OSOM 37 ± 1%, and mTAL of ISOM 46 ± 7%, respectively (**Figure 1D**). Mosaic ROMK signal was comparable across species using mouse and rat kidney samples with similar numerical outcome and no significant sex difference (**Supplemental Figure 1B-D**). In contrast to the mosaicism of ROMK protein signal, *Kcnj1* mRNA encoding for ROMK, was present in all TAL cells as detected by *in situ* hybridization (**Supplemental Figure 2**).

**Figure 1.**
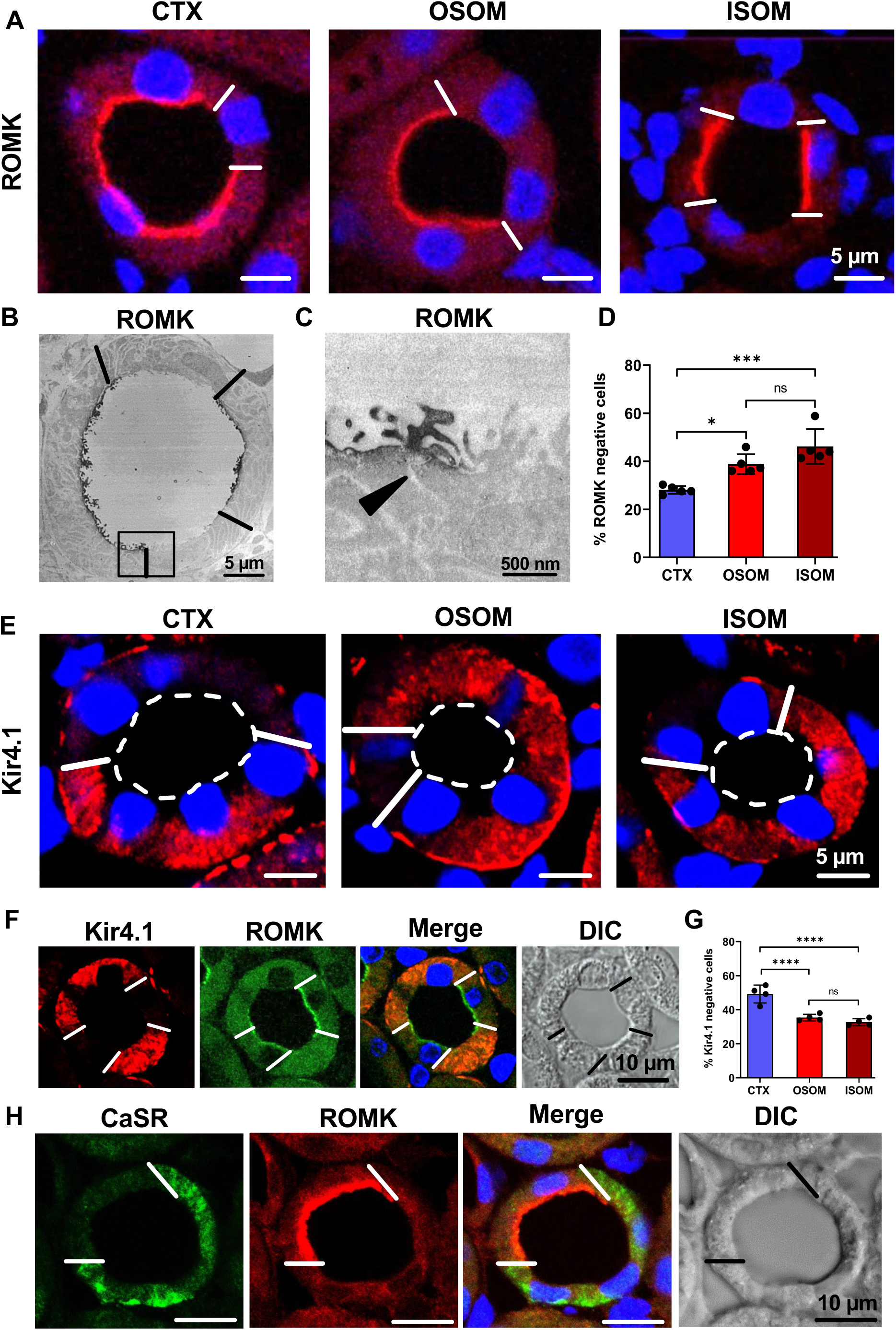
Immunostaining for ROMK, Kir4.1, and CaSR distribution in rat kidney TAL. **(A)** ROMK immunostaining is identified by luminal signal in a subset of cells lining TAL profiles across renal parenchymal zones; DAPI blue nuclear staining. White bars in epithelia identify cell borders between ROMK-positive and ROMK-negative cells. **(B,C)** Transmission electron microscopy showing overview **(B)** and detail **(C)** of ROMK-positive and ROMK-negative adjacent cells; arrowhead marks junctional area between positive and negative cells **(C)**; immunoperoxidase staining. Scale bars indicate magnification. (**D)** Numerical evaluation of ROMK-negative cells across the zones; values are means ± SD from n=5 rats; * *P* < 0.05; *** *P* < 0.001; ns, not significant by ANOVA and post-hoc comparisons. **(E)** Kir4.1 immunostaining is identified as basolateral signal in a subset of medullary (m) TAL epithelial cells. **(F)** Double immunostaining for Kir4.1 and ROMK shows mutually exclusive signals in mTAL. **(G)** Numerical evaluation of Kir4.1-negative cells across zones; values are means ± SD from n=4 rats; **** *P* < 0.0001; ns, not significant by ANOVA and post-hoc comparisons. **(H)** Double immunostaining for CaSR and ROMK show mutually exclusive signals in mTAL. White bars mark the borders between ROMK-positive and -negative cells. To reflect underlying structure, a differential interference contrast (DIC) filter was used. DAPI blue nuclear staining; scale bars indicate magnification.

The Kir4.1 inwardly rectifying K^+^ channel, known to be present along the basolateral membrane of TAL cells (21), revealed mosaicism as well that was mutually exclusive to ROMK expression in TAL across zones. Commonly, ROMK-negative TAL cells thus expressed significant Kir4.1 signal, whereas ROMK-positive cells were devoid of Kir4.1 signal (**Figure 1E and F**). Kir4.1-negative cells amounted to 48 ± 8% in cTAL, 36 ± 6% in mTAL of OSOM, and 33 ± 7% in mTAL of ISOM (**Figure 1G**). Further mosaicism was detected for the CaSR with strong signal in cTAL and gradually decreasing strength in mTAL. Cells expressing either CaSR or ROMK were typically found in mTAL, whereas in cTAL, ROMK-positive cells commonly also expressed CaSR (**Figure 1H**).

Despite the uniform NKCC2 signal in TAL, antibody staining of phospho-S130 NKCC2 (pNKCC2), which reflects the activated state of the cotransporter (22), displayed mosaicism in rat kidney TAL as well. The distribution of pNKCC2 signal was inversely correlated with ROMK in mTAL while in cTAL, pNKCC2 signal was stronger than in mTAL and displayed less mosaicism. Numerically, 58 ± 11% of mTAL cells were pNKCC2-negative TAL in OSOM, 57 ± 6% in ISOM, while only 5 ± 7% in cTAL (**Supplemental Figure 1E and F**).

### Potassium Channels In Macula densa

In macula densa, ROMK-signal was evenly or irregularly distributed (**Supplemental Figure 3**). In the post-macula segment, ROMK distribution did not differ from that in upstream cTAL. ROMK-negative cells were mostly solitary, but two adjacent negative cells were also found (**Supplemental Figure 3**).

### ROMK Is Expressed In Cells With Lower Total Potassium Conductance

The immunolocalization studies suggest differential K^+^-conductive pathways among TAL cells, and as two types of TAL cell conductive properties were reported in hamster (11), we performed whole-cell voltage-clamp experiments in isolated and split-open mouse cTAL. **Figure 2A** shows examples of patch-clamp recordings from tertiapin-Q (TPNQ, 400 nM)-sensitive K^+^ currents indicating ROMK at −40 mV (23) and residual Ba^2+^-sensitive K^+^ currents indicating K^+^ currents other than ROMK. **Figure 2B** summarizes the results, showing that there are two types of cTAL cells, one displaying TPNQ-sensitive ROMK channel activity (390 ± 42 pA; six of fourteen cells) and another with no ROMK channel activity (eight of fourteen cells); the latter display higher Ba^2+^-sensitive K^+^ currents compared to the former. As whole-cell electrophysiology does not permit luminal and basolateral membranes to be perturbed individually, we perfused dissected cTAL segments. Membrane voltage changes were measured by confocal microscopy using the membrane voltage dye Di-8-ANEPPS (**Figure 2C**). At constant luminal K^+^ concentration, the basolateral K^+^ concentration was increased to depolarize cells that have a high fractional K^+^ conductance at the basolateral membrane. During this maneuver, some TAL cells depolarized, whereas others did not.

**Figure 2.**
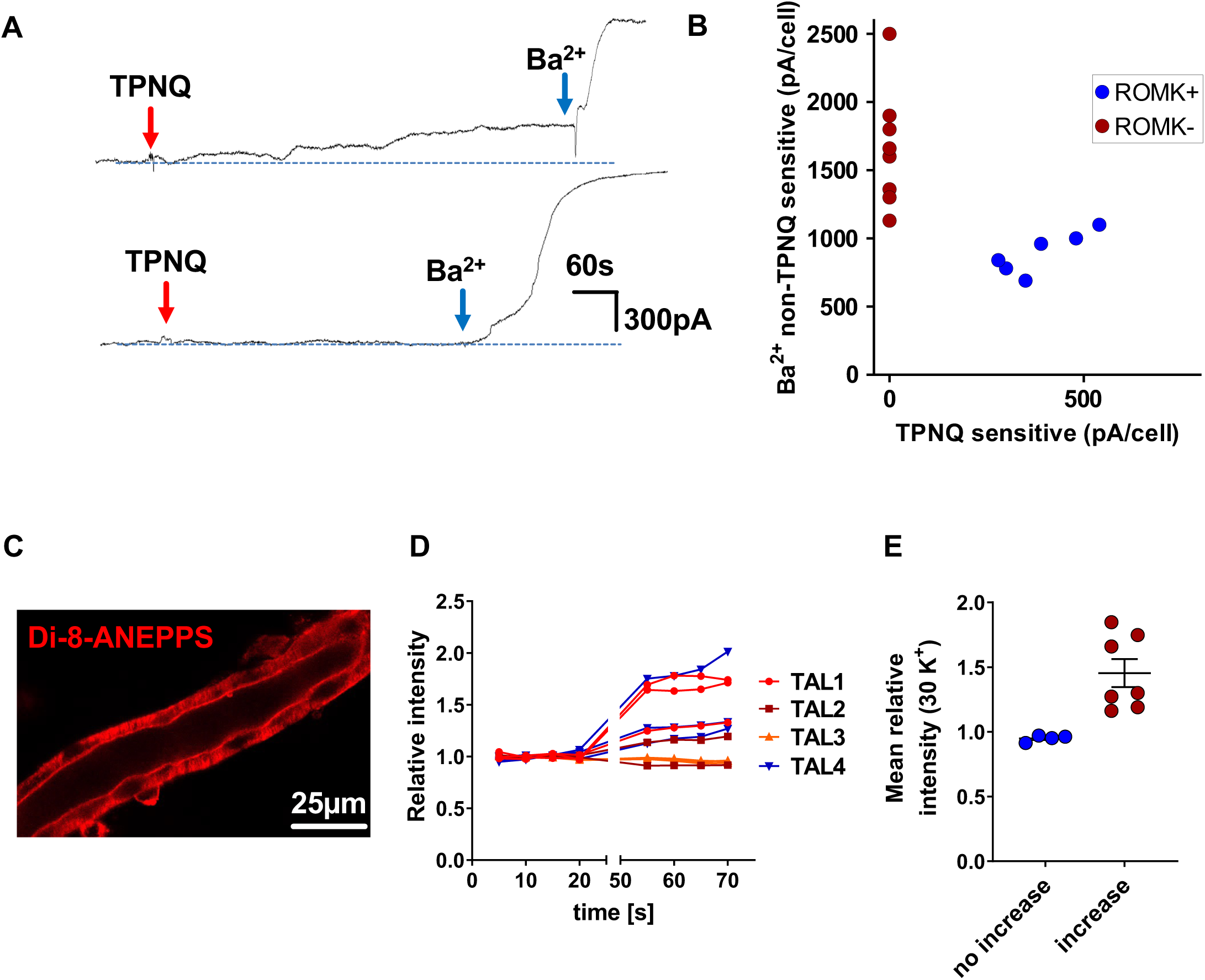
Functional evidence for ROMK activity in cells with lower total potassium conductance. **(A)** Whole-cell recordings show tertiapin-Q (TPNQ)-sensitive K currents at −40 mV and Barium-sensitive K^+^ currents at the same voltage in two representative TAL cells. Red arrows indicate addition of 400 nM TPNQ to the bath; blue arrows indicate addition of 1 mM Ba^2+^ to the bath. **(B)** Dot plot summary of experiments of TPNQ-sensitive and Ba^2+^-sensitive K^+^ currents at −40 mV with whole cell recording. There are two types of cTAL cells, one (six out of fourteen cells) with TPNQ-sensitive ROMK channel activity (390 ± 42 pA), the other (eight out of fourteen cells) with no ROMK channel activity. Note that ROMK-negative cells have a higher Ba^2+^-sensitive K^+^ currents than ROMK-positive cells. Results were obtained from 14 experiments (tubules). **(C)** Confocal image of an isolated perfused TAL after Di-8-ANEPPS loading. Fluorescence intensity corresponds to membrane voltage and dye concentration. **(D)** Relative intensity of 11 individual cells from 4 TAL tubules under control (3.6 mmol/l) or high basolateral K^+^ concentration (30 mmol/l). **(E)** Mean relative intensities in the presence of high basolateral K^+^ concentration suggesting two distinct cell types.

Relative fluorescence intensities before and after exposure to basolateral 30 mmol/l K^+^ are shown in **Figure 2D** from 11 individual cells of 4 different TAL tubules. Fluorescence increase indicates depolarization in the presence of 30 mmol/l K^+^ at the basolateral side which is indicative of a basolateral K^+^ conductance. No response indicates that the membrane voltage completely depends on luminal K^+^ channels as the luminal K^+^ concentration remained constant throughout the experiment; two distinct cell types could thus be identified (**Figure 2E**).

### Immunohistochemistry Links Claudin Mosaicism with ROMK

Mosaicism was further confirmed in the expression of the two key tight junctional proteins, Cldn10 and Cldn16, along TAL as previously reported (15, 24). Accordingly, Cldn10-positive cells were encountered along the entire cTAL and mTAL, whereas Cldn16-positive cells were restricted to TAL within cortex, OSOM, and outermost ISOM (**Figure 3**). At the cellular level, Cldn10 immunoreactivity decorated the basolateral membrane and tight junction, as reported (25), whereas Cldn16 was strictly confined to the junctional location. In cortex and OSOM, ROMK-negative cells were entirely surrounded by a Cldn16-positive junctional belt, whereas ROMK-positive cells revealed two different types of junctional belt composition. In one, ROMK-positive cells surrounded by other ROMK-positive cells expressed a Cldn10 junctional belt at their entire circumference. In the other, ROMK-positive cells bordering ROMK-negative cells displayed Cldn16 junctional belt where contacting each other, so that ROMK-positive cells, but not ROMK-negative/Kir4.1-positive cells revealed a heterogeneity between Cldn10 and Cldn16 signals along their junctional portions (**Figure 3A and B, diagrammed in 3C**).

**Figure 3.**
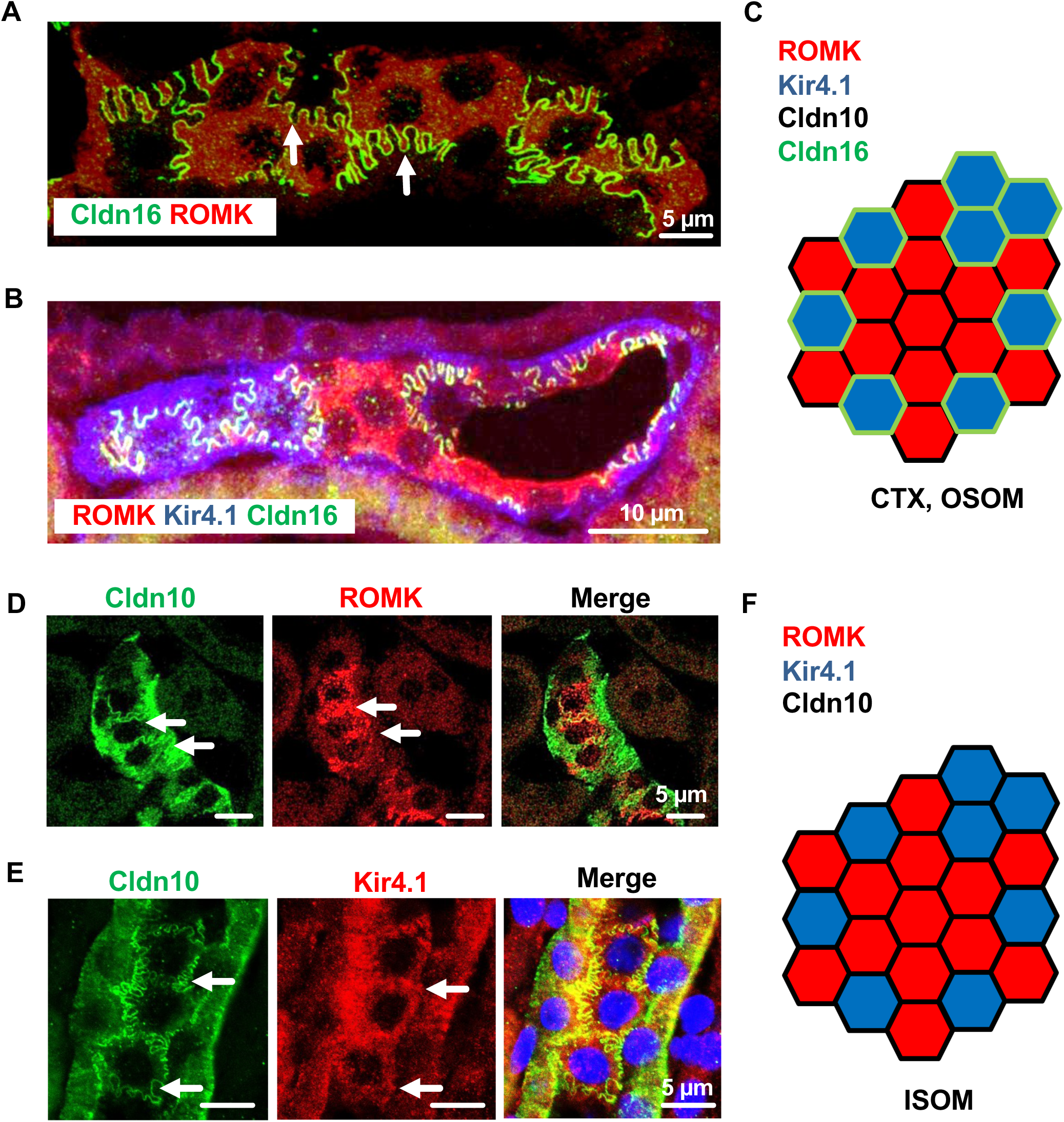
Immunostaining for claudin 10, claudin 16, ROMK and Kir4.1 in rat kidney. Images were taken in subluminal tangentional plane of the tubules**. (A)** Double immunostaining for Claudin (Cldn)16 and ROMK in mTAL of OSOM; note absence of Cldn16 signal between ROMK-positive cells, but its presence when lining ROMK-negative cells (arrows). **(B)** Triple immunostaining staining for ROMK, Kir4.1 and Cldn16 in cTAL; red and purple signals distinguish ROMK-positive from ROMK-negative/Kir4.1-positive cells, respectively. Note absence of Cldn16 from junctions between ROMK-positive cells, but its presence between ROMK-negative/Kir4.1-positive cells. **(C)** Schematic drawing demonstrates the differences with respect to ROMK-positive, Kir4.1-negative as well as ROMK-negative, Kir4.1-positive cells and their Cldn10- or Cldn16-specific junctional lining in TAL of cortex and OSOM. **(D)** Double immunostaining for Cldn10 and ROMK in mTAL of ISOM; junctional Cldn10 signals (arrows) between ROMK-positive cells. **(E)** Double immunostaining for Cldn10 and Kir4.1 in mTAL of ISOM; tight junctions of Kir4.1-positive cells express Cldn10. **(F)** Schematic drawing demonstrates ROMK-positive, Kir4.1-negative as well as ROMK-negative, Kir4.1-positive cells and their Cldn10-specific junctional lining in TAL of ISOM. Scale bars indicate magnification.

These junctional staining patterns were confirmed in mouse isolated cTAL segments (**Supplemental Figure 4**), where mixing of Cldn10 and Cldn16 signals in a single tight junction was not observed. Some features in the ISOM were different. As noted above, cells in this region express only Cldn10 and this was true for both ROMK-positive and ROMK-negative cells. As in cortex and OSOM, cells expressing Kir 4.1 did not have apical ROMK (**Figure 3D and E, diagrammed in 3F**).

### Enriched Single-Nucleus RNA Sequencing Reveals Distinct TAL Cell Types

To classify TAL cell types and complement the anatomical and functional studies above, we enriched TAL nuclei for single-nucleus (sn) RNA sequencing. Given that TAL cells constitute a small fraction of kidney tissue and exhibit heterogeneity across and within kidney zones, we employed an enrichment approach to enhance resolution. First, we generated NKCC2-INTACT mice by crossing NKCC2-Cre and CAG-Sun1/sfGFP lines, enabling inducible TAL-specific nuclear labeling, as we previously described for DCT snRNA-Seq (26). Tamoxifen-induced recombination efficiently targeted TAL cells (**Figure 4A**).

**Figure 4.**
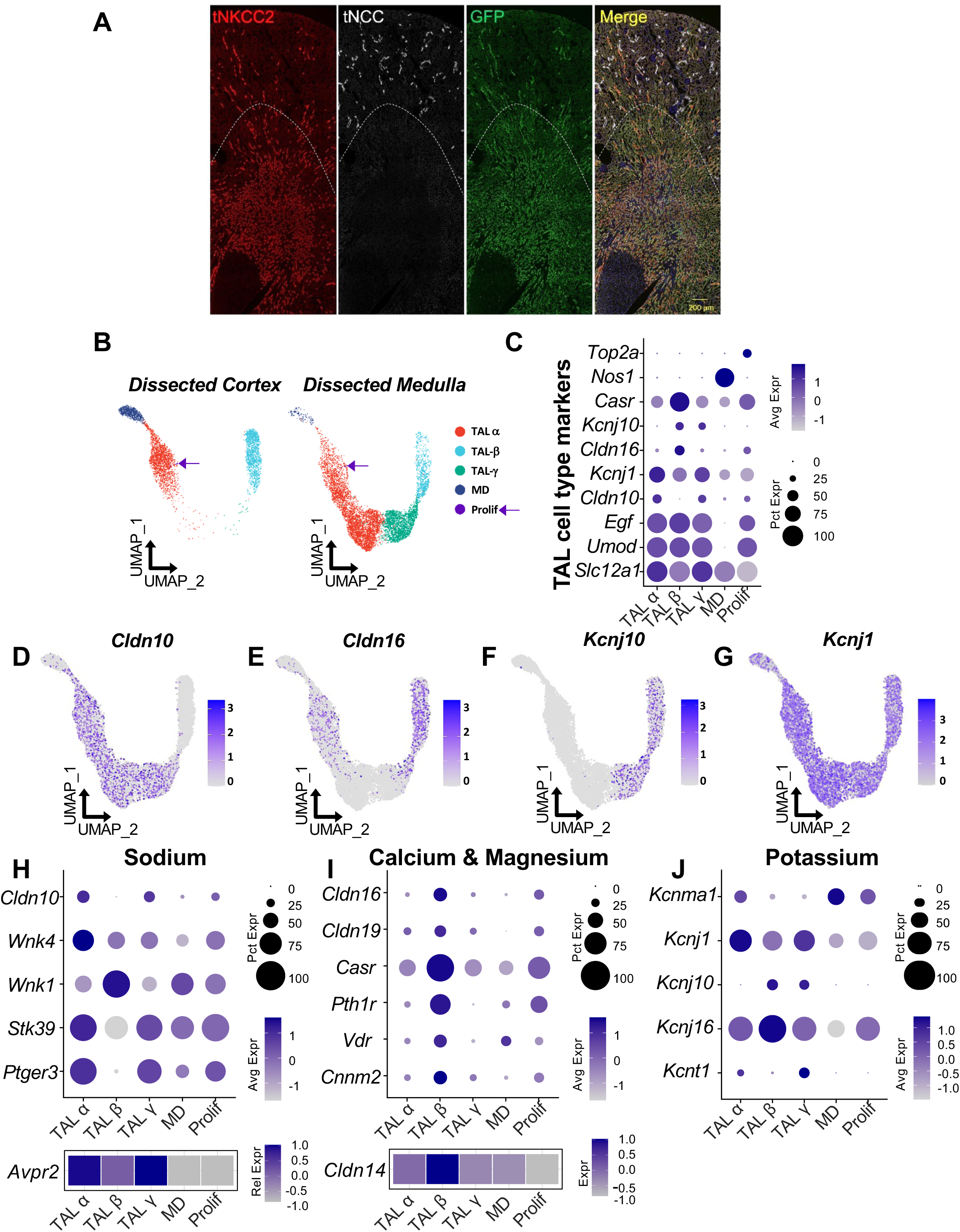
TAL cell diversity revealed by enriched single-nuclei RNA-seq analysis in the mouse kidney. (**A**) Immunofluorescence staining of NKCC2-INTACT mouse kidneys shows tNKCC2 (red), tNCC (white), and GFP (green). Tamoxifen-induced recombination specifically labels NKCC2+ cells, with GFP-positive nuclei colocalizing exclusively in NKCC2-expressing cells. (**B**) Uniform manifold approximation and projection (UMAP) projections of enriched single-nuclei RNA-seq data show TAL cell populations split by kidney region (cortex and medulla). Identified clusters include TAL-α (Claudin-10–positive, Kir4.1–negative), TAL-β (Claudin-16–positive, Kir4.1–positive), TAL-γ (Claudin-10–positive, Kir4.1–positive), macula densa (MD; Nos1–positive), and proliferating (Prolif; Top2a–positive) cells. (**C**) Dot plot displaying key TAL cell-type markers, including *Slc12a1*, *Cldn10*, *Cldn16*, *Kcnj10*, *Kcnj1*, *Nos1*, *Top2a.* UMAP projections displaying the expression of *Cldn10* (**D**), *Cldn16* (**E**), *Kcnj10* (**F**), and *Kcnj1* (**G**) genes. Dot plots and heatmaps illustrate gene expression patterns associated with: (**H**) Sodium transport (*Cldn10*, *Wnk4*, *Wnk1*, *Stk39*, *Ptger3*, *Avpr2*), which are enriched in TAL-α and TAL-γ clusters. (**I**) Calcium and magnesium transport (**C** *ldn16*, *Cldn19*, *Casr*, *Pth1r*, *Vdr*, *Cnnm2*, *Cldn14*), with higher expression in TAL-β. (**J**) Potassium transport (*Kcnma1*, *Kcnj1*, *Kcnj10*, *Kcnj16*, *Kcnt1*). Data normalization and scaling: Gene expression data were normalized and z-score scaled to enable comparison of relative expression levels across clusters. “Avg Exp” (Average Expression) and “Exp” (Expression) represents the z-scored mean gene expression within a cluster, while “Pct Exp” (Percent Expressed) denotes the percentage of cells in a cluster with detectable expression of the gene.

Transcriptomic analysis of enriched NKCC2-INTACT nuclei showed uniform *Slc12a1* (NKCC2) expression, validating TAL cell identity (**Supplemental Figure 5A**). Additionally, kidneys were manually dissected into cortical and medullary regions prior to sequencing, with UMAP visualization revealing clear segregation of TAL cells between these regions (**Supplemental Figure 5B and C**).

Results identified three major and two minor TAL cell clusters (**Figure 4B**). The major clusters were classified as TAL-α (Claudin 10-positive, Kir4.1-negative), TAL-β (Claudin 16– positive, Kir4.1-positive), and TAL-γ (Claudin 10-positive, Kir4.1-positive) (**Figure 4C, Supplemental Table 1**). TAL-α and TAL-β cells were present in both cortical and medullary regions, whereas TAL-γ cells were exclusive to the medulla (Figure 4B). Among the minor clusters, one exhibited *Nos1* expression, indicating that it represents macula densa cells, while the other expressed *Top2a*, representing a small population of proliferating cells (**Figure 4C**). These findings align with prior transcriptomic analyses of cortex (17, 18) but provide enhanced resolution, particularly within the medullary regions.

The enriched TAL dataset revealed previously unrecognized molecular heterogeneity, particularly within the medullary TAL. Notably, *Kcnj10* (Kir4.1) showed distinct patterns: in the cortex, it was exclusively expressed in Claudin 16-positive (TAL-β) cells, while in the medulla, it was expressed in both Claudin 16-positive (TAL-β) and Claudin 10-positive (TAL-γ) cells (**Figure 4B, D, E, F, Supplemental Table 1**). Present immunostaining and previous studies (28) suggest that medullary TAL-β cells reside in the OSOM, whereas TAL-γ cells are likely restricted to the ISOM.

Although *Kcnj1* (ROMK) transcripts were detected across all clusters, as noted above (**Supplemental Figure 2**) they were significantly enriched in Kir4.1-negative TAL-α cells, corresponding to apical ROMK expression (**Figure 4G**). Notably, complete cell type identification requires apical ROMK localization, although transcriptomic data alone can be used to identify TAL-α, TAL-β, and TAL-γ cells (**Supplemental Table 1**).

Differentially expressed gene (DEG) analysis further revealed functional distinctions among TAL subtypes. TAL-α and TAL-γ cells (Claudin 10–positive) exhibited higher expression of genes associated with sodium transport (*WNK4*, *Stk39* [SPAK], *Avpr2*), as well as arachidonic acid signaling (*Ptger3*), and the organic anion transmembrane transporter Slco3a1 (Oatp3a1), which has been implicated in prostaglandin transport (27) (**Figure 4H, Supplemental Figure 6A**). In contrast, TAL-β cells (Claudin 16–positive) showed enriched expression of genes involved in calcium and magnesium homeostasis, including Claudin 19, Claudin 14, *Casr*, the parathyroid hormone receptor (*Pth1r*), the vitamin D receptor (*Vdr*), and Cyclin and CBS domain divalent metal cation transport mediator 2 (*Cnnm2*) (**Figure 4I**). Notably, WNK1 expression was enriched in TAL-β cells, whereas WNK4 was more highly expressed in TAL-α cells (**Figure 4H**).

Other potassium channels also exhibited differential expression. *Kcnma1*, which encodes the Potassium Calcium-Activated Channel Subfamily M Alpha 1 (BK channel) was predominantly expressed in TAL-α cells, while *Kcnj16* (Kir5.1), a channel that forms heterotetramers with Kir4.1, was widely expressed across TAL types but with highest expression in TAL-β cells (**Figure 4J**). *Kcnt1*, recently linked to the 70 pS apical potassium channel along TAL (28), was exclusive to medullary TAL-α and TAL-γ cells (**Figure 5A**, **Supplemental Figure 5F**). A summary of the top 10 DEGs for each TAL cell type is presented in **Supplemental Figure 5G**, with a comprehensive list of DEGs included in the supplemental material. Additional highlighted genes are presented in Supplemental Figure 6.

**Figure 5.**
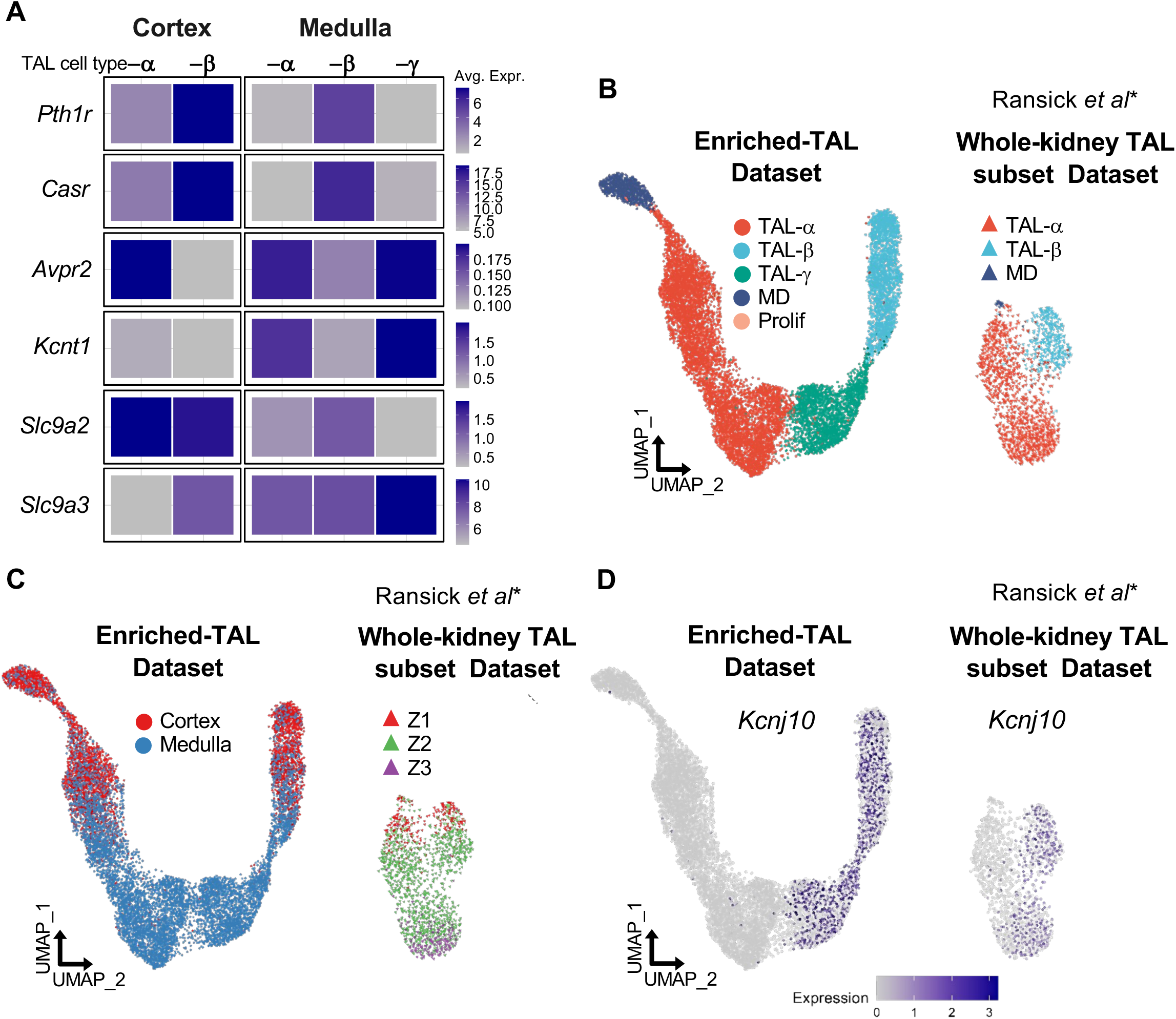
Regional Expression of Key Proteins in TAL Cell Types and Comparative Analysis of TAL Diversity: Enriched-TAL vs Whole-Kidney TAL Subset in RNA-seq Datasets Heatmap showing regional gene expression patterns in TAL cell types. Pth1r and Casr transcripts are highest in TAL-β cells, peaking in the cortex. Avpr2 and Kcnt1 are enriched in Claudin 10–positive TAL-α and TAL-γ cells, with highest levels in the medulla. Slc9a2 (NHE2) is primarily expressed in the cortex (TAL-α and TAL-β), while Slc9a3 (NHE3) is medulla-enriched, peaking in TAL-γ cells. Data normalization and scaling: Gene expression data were normalized and scaled using a z-score to compare relative expression levels across clusters. “Avg Exp” (Average Expression) indicates the z-scored mean gene expression within a cluster. (**B-D**) Comparative analysis of the enriched-TAL dataset with *Slc12a1*-positive TAL subsets extracted from a previously published whole-kidney RNA-seq dataset (16). (**B**) UMAP projection comparing TAL cell types. The enriched-TAL dataset reveals three distinct TAL cell types: TAL-α, TAL-β, and TAL-γ, alongside macula densa (MD) and proliferating (Prolif) cells. In contrast, the whole-kidney TAL subset (16) predominantly identifies two TAL cell types (TAL-α and TAL-β) and MD cells. (**C**) UMAP projection illustrating the origin of TAL cells by kidney zone according to authors. In the enriched-TAL dataset, cells are derived from dissected cortex and medulla. In the whole-kidney dataset, kidney zones are designated as Z1 (cortex), Z2 (outer medulla), and Z3 (inner medulla) (16). (**D**) UMAP projection displaying Kcnj10 (Kir4.1) expression in both the enriched-TAL dataset and the whole-kidney TAL subset (16).

### Regional Expression Patterns and Functional Insights

Our combined molecular and hand dissection approaches provided new insight into regional gene expression patterns. **Figure 5A** compares relative expression across cell types and regions. *Pth1r* and *Casr* transcripts were most abundant in TAL-β cells, with peak expression in the cortex. Interestingly, a subset of cortical Claudin 10–positive cells co-expressed *Pth1r* and *Casr*. In contrast, *Avpr2* and *Kcnt1* transcripts were enriched in Claudin 10–positive TAL-α and TAL-γ cells, with highest levels in the medullary regions. Consistent with prior studies (14), *Slc9a2* (NHE2) was predominantly expressed in the cortex, with similar levels in TAL-α and TAL-β cells, whereas *Slc9a3* (NHE3) was enriched in the medulla, showing peak expression in TAL-γ cells.

As many published datasets do not contain anatomical information, we used the top 10 DEGs genes from cortical and medullary TAL cells to generate a “cortex score” and a “medullary score” (**Supplemental Figure 7**), serving as a robust spatial marker for TAL regions. Additional relevant DEGs are provided in **Supplemental Figure 7**.

### TAL Cell-Types and Anatomical Location Are Conserved Across Species

Immunolocalization results indicated similar ROMK expression heterogeneity in mouse, rat, and human (**Supplemental Figure 1**). To extend this, we examined TAL cell types across species, using published datasets from mouse, human, and rat kidneys (**Supplemental Figure 8-10**). The enhanced resolution obtained using the INTACT method was clearly evident when we compared our data with data from another mouse dataset (16) (**Figure 5B-D**). There, we identified TAL-α cells, characterized by high Cldn10 expression, and TAL-β, marked by high Cldn16 expression (**Supplemental Figure 8B**), but TAL-γ cells did not form a distinct cluster (**Supplemental Figure 8C-E**).

Analysis of the Kidney Precision Medicine Project (KPMP) human dataset (29) also revealed TAL-α and TAL-β clusters (**Supplemental Figure 9A-D**). As with other datasets lacking TAL enrichment, TAL-γ cells were rare. Nevertheless, there was a strong and significant correlation between mouse and human TAL-α and TAL-β differentially expressed genes (DEGs) (**Supplemental Figure 9H-I**).

In rats, unsupervised clustering of TAL cells also revealed clusters based on Claudin 10 and Claudin 16 expression. In this dataset, TAL-α cells further segregated into two subtypes (**Supplemental Figure 10A-D**), likely from distinct kidney regions, but these cells could not be identified as TAL-γ. Regional markers (*Clcnkb*, *Enox1* for the cortex; *Clcnka*, *Slc4a7*, *Ank2* for the medulla) confirmed these spatial distinctions (**Supplemental Figure 10E**).

Across species, we observed that Claudin 10–positive cells (TAL-α cells) consistently express key factors involved in sodium transport, as well as arachidonic acid signaling genes (**Supplemental Figure 8F, 9E, 10G**) whereas, Claudin 16–positive (TAL-β) cells showed higher expression of genes associated with calcium and magnesium homeostasis (**Supplemental Figure 8G, 9F, 10H**). DEG profiles of TAL-α and TAL-β cells were highly correlated between species (**Supplemental Figures 9H-I and 10J-K**). A comprehensive list of DEGs for TAL-α and TAL-β cells across species is provided in the supplementary material.

Furthermore, sex-based differences in TAL cell proportions were observed, with females showing a higher percentage of TAL-β cells in both mice and humans (**Supplemental Figures 8J and 9J**).

### Ultrastructural Features of TAL

The results described above indicate the existence of 3 cell types along the TAL, differing importantly in the expression of K^+^ channels. Some have speculated that differences in TAL conductance properties correspond to the appearance of “rough” and “smooth” TAL cells (11, 30). To examine this, we studied the morphology of medullary and cortical TAL epithelia under basal conditions using transmission (TEM) and scanning electron microscopy (SEM) in rat and mouse. In rat, TEM of ISOM showed that the luminal cell aspect of mTAL cells was a smooth plateau with few scattered, stubby microvilli, whereas cell borders were densely packed with marginal microvilli. Towards outer stripe and cortex, meandering of the TAL cell borders and density of marginal microvilli increased progressively along with the number of microvilli of the luminal plateau (**Figure 6 A-I**). Cells characteristically revealed a subapical vesicular compartment throughout with tightly packed vesicles. Subapical vesicles were most abundant in ISOM, less so in OSOM, and scarce in cTAL (**Figure 6 D-F**), confirming earlier data (31). Despite clearly visible axial heterogeneity of TAL epithelia in the respective kidney parenchymal zones, no evidence for luminal cell surface heterogeneity was detected within a given zone; this was confirmed by SEM showing no mosaic distribution of “smooth (S)” or “rough (R)” cells in contrast to earlier descriptions (11, 32) (**Figure 6 J-L**). Similar findings were obtained in mouse kidney (**Figure 6M**).

**Figure 6.**
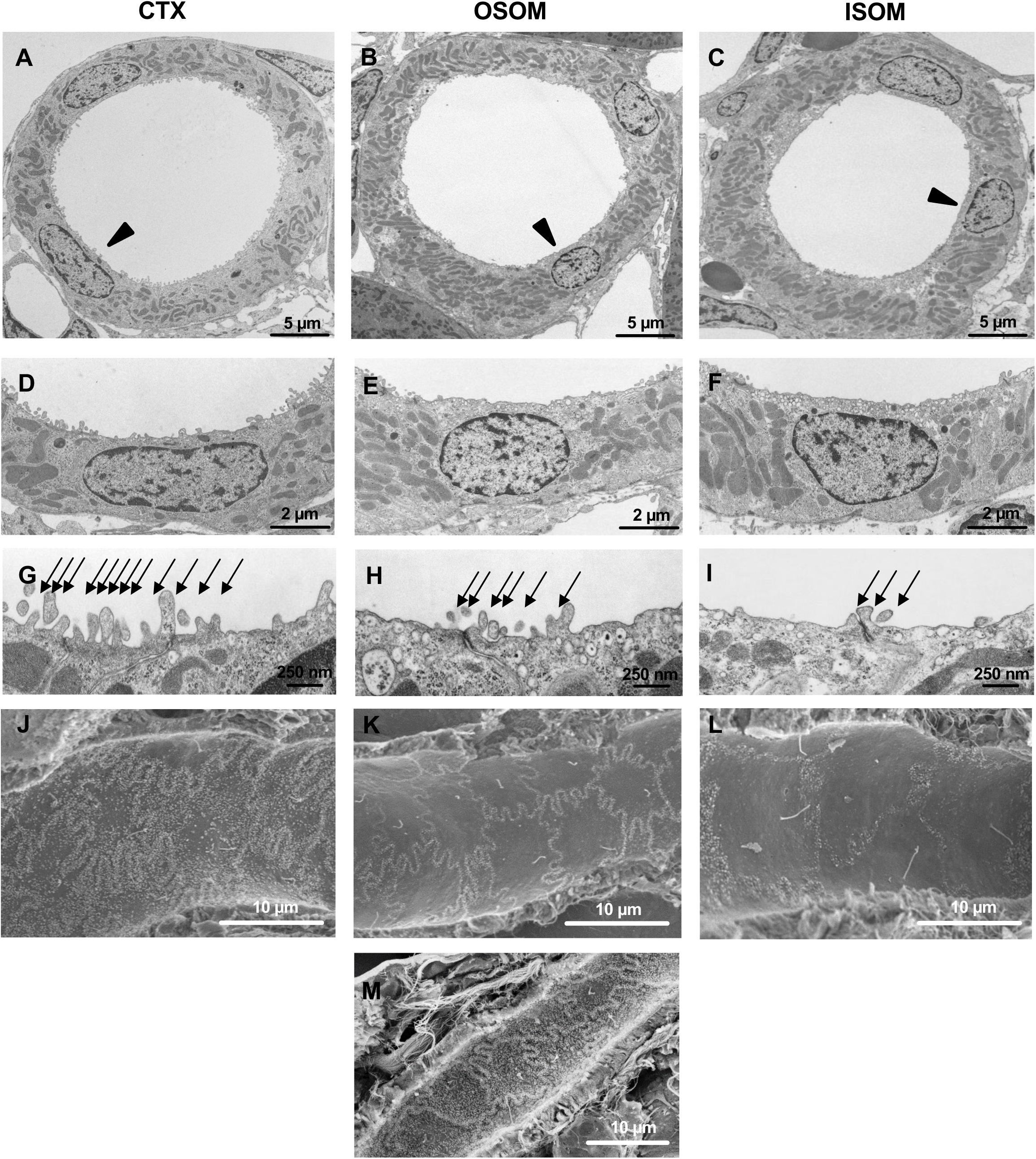
Ultrastructural landmarks in TAL, rat and mouse kidney. TAL from cortex (CTX), OSOM, and ISOM. **(A-F)** Cross sectional profiles **(A-C)** with increasing cell height towards ISOM; arrowheads indicate sites of higher magnification **(D-F)** by transmission electron microscopy. Note minor differences in microvillar density along the course of TAL. **(G-H)** Tight junctional fields show numbers of luminal microvilli along luminal cell borders decreasing from CTX to ISM (arrows). **(J-M)** Scanning electron micrographs showing luminal cell aspects of TAL epithelium in rat **(J-L)** and mouse kidney **(M)**. Note decreasing complexity of cell borders and numbers of microvilli in rat as well as general absence of cell heterogeneity. Scale bars indicate magnification.

### Summary of Results

Results have been schematized with respect to cell type characterization (**Figure 7 and Supplemental Figure 11**)

**Figure 7.**
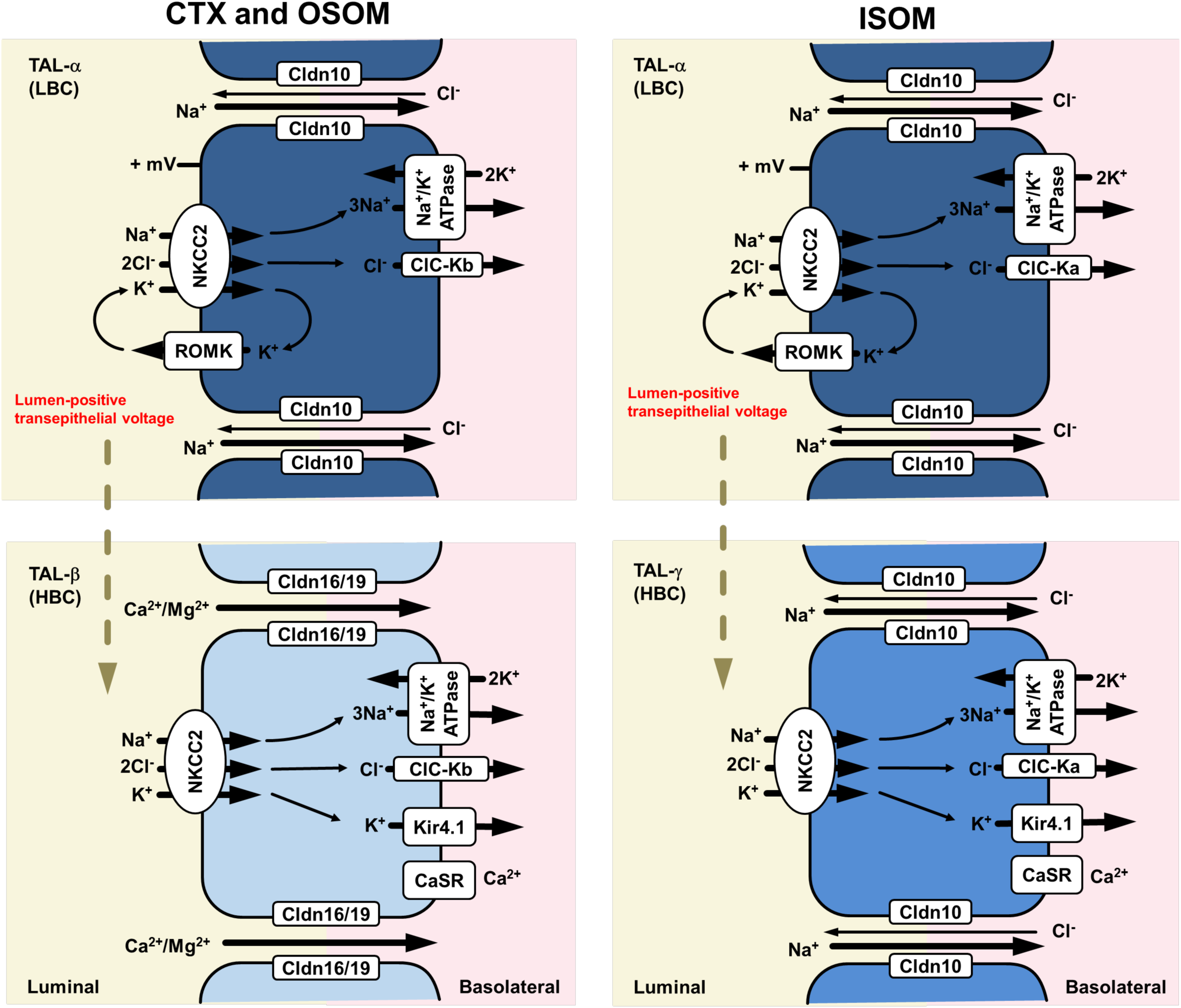
Schematic drawing on cell type heterogeneity along TAL. Mechanistic model of TAL cellular heterogeneity. All cells express luminal NKCC2, basolateral ClC channels and Na^+^,K^+^-ATPase to reabsorb NaCl via the transcellular path. TAL-α cells of all zones are equipped for paracellular Na^+^ absorption via Cldn10. They express luminal ROMK and show low basolateral K^+^ conductance (LBC) to generate a lumen-positive voltage. TAL-β cells of CTX and OSOM comprise the equipment for paracellular Ca^2+^/Mg^2+^ reabsorption via Cldn16/19. They lack luminal ROMK and show high basolateral K^+^ conductance (HBC). TAL-γ cells of ISOM are equipped for paracellular Na^+^ absorption via Cldn10; they lack luminal ROMK and show high basolateral K^+^ conductance (HBC). TAL-β and TAL-γ cells per se do not contribute to transepithelial voltage, but adjacent LBC type TAL-α cells still provide the electrotonic driving force (dotted arrow) to support K^+^ reabsorption and paracellular passage of Na^+^ (TAL-α & TAL-γ cells) or Ca^2+^/Mg^2+^ (TAL-β cells). Data are from previous work and current results.

## Discussion

The TAL reabsorbs substantial amounts of filtered sodium, chloride, potassium, calcium, and magnesium, and plays a central role in urinary concentration. It must be able to modulate ion transport selectively to meet homeostatic needs. Axial heterogeneity along the TAL is well recognized, but nearly all TAL cell models suggest a single TAL cell type, albeit some with variants (33). The present results show (**Figure 7**) that the mammalian TAL comprises three distinct cell types in addition to macula densa cells; each cell type expresses unique transport-related genes and regulatory proteins, suggesting that they respond to different stimuli and play unique roles in homeostasis. Combining immunolocalization, enriched snRNA-Seq, and functional analyses, we now provide a new model for solute transport pathways along the TAL that accounts for several previously confusing findings.

The results show that TAL cells express either apical ROMK or basolateral Kir4.1, not both, and that cells lacking apical ROMK have higher total K^+^-conductance. These results are consistent with early work showing that hamster TAL cells and amphiuma early distal tubule (its diluting segment) cells exhibit two patterns of membrane conductance labeled as HBC and LBC (10, 11). Additionally, the results show that cells with apical ROMK (TAL-α cells) express the Na^+^-permeable claudin 10 at their tight junction, allowing them to both generate a transepithelial voltage and transport Na^+^ efficiently both through and around cells. As they also express NKCC2 at the apical membrane, TAL-α cells correspond to the traditional model for TAL cells in which nearly all K^+^ that enters across the apical membrane is recycled (34, 35). As might be expected, these claudin 10-expressing cells differentially express genes related to Na^+^ transport and water balance, such as *Stk39* (encoding SPAK) (36), *Avpr2* (encoding the vasopressin receptor), and *Ptger3* (encoding prostaglandin E3 receptor) (37).

In contrast, cells in the cortex and OSOM that express Kir4.1 at their basolateral membrane also have NKCC2 at the apical membrane but do not exhibit apical ROMK; these cells express claudin 16 (TAL-β cells), marking them as Ca^2+^ and Mg^2+^ transporting cells. As might be expected, these cells differentially express genes encoding the PTH and vitamin D receptors. The absence of apical ROMK means that this cell type (TAL-β cells) does not contribute directly to transepithelial voltage generation. Yet, as Ca^2+^ and Mg^2+^ reabsorption takes place via divalent ion-permeable claudin 16 (and 19), it must be driven by the voltage generated by adjacent TAL-α cells. Additionally, in the cortex, where substantial divalent ion reabsorption occurs, the transepithelial voltage rises further, owing to a Na^+^ diffusion potential across Na+ selective TAL-α cell tight junctions, reaching as high as 30 mV (33). In the ISOM, there is a third cell type that, like TAL-β cells, expresses Kir4.1 at the basolateral membrane and lacks ROMK at the apical membrane. These cells, however, have claudin 10 rather than claudin 16 within their tight junctions. We have labelled these TAL-γ cells, as they differ from the others, but these cells are often missed in analyses that don’t enrich for TAL cells.

A key current finding is that ROMK/Kir4.1 mosaicism is present all along the TAL. Ultrastructural analysis clearly identified the onset of the ROMK signal at the tight junctions between ROMK-negative and ROMK-positive cells, confirming an earlier suggestion that this is not an artifact (12, 13). Our electrophysiological findings in mice align with the transcriptomic and immunolocalization results, thereby extending prior transcriptomic studies in other species that often suggested two functional cell types, identified by differential claudin expression (10, 11). By integrating highly enriched transcriptomic analysis with immunolocalization and electrophysiology, and focusing on TAL cells across all kidney regions, a clear cell type differentiation appears. Wang and colleagues identified Kir4.1 (with Kir5.1) as responsible for a K^+^ conductive pathways at the basolateral membrane of cTAL cells. They reported that 40 pS K^+^ channels were absent in the basolateral membrane of TAL cells from mice lacking *Kcnj10* (Kir4.1), indicating their identity (21). Interestingly, in light of the current results, they noted by that Kir4.1 staining was ‘*not uniform*’ along the TAL. Here, we used the ROMK blocker TPNQ and the nonspecific K^+^ channel blocker barium and found that only a fraction of TAL cells exhibit ROMK activity (TPNQ-sensitivity), consistent with our immunocytochemical results. These experiments also indicate that cells lacking ROMK exhibit higher whole cell Ba^2+^-sensitive K^+^ conductance, again aligning with the mosaicism of K^+^ channels described (10, 11). Further, in perfused tubules, where we could control apical and basolateral composition selectively, we found that only a fraction of cells respond to changes in basolateral [K^+^]. Showing that this basolateral conductance is provided by cell-specific expression of Kir4.1 will require additional studies. The loop of Henle is known to reabsorb K^+^, but mechanisms have been unclear.

Although some K^+^ is believed to traverse the paracellular pathway driven by the transepithelial voltage, transcellular absorption also occurs (38). The reciprocity of K^+^ channels among different cell types is likely essential for TAL K^+^ reabsorption. TAL models throughout the years show that nearly all the K^+^ traversing the apical membrane via NKCC2 recycles back into the lumen (39, 40). Thus, owing to their high apical and low basolateral conductance, TAL-α cells do not mediate transcellular K^+^ reabsorption. Our data argue that this K^+^ reabsorption is mediated by TAL-β and TAL-γ cells, which have NKCC2, lack an apical K^+^ exit pathway, and have high basolateral K^+^ conductance. Therefore, as shown in **Figure 7**, K^+^ entering TAL-β and TAL-γ cells must leave via the basolateral membrane, leading to net K^+^ reabsorption. The distinction between K^+^ reabsorbing cells (TAL-β and TAL-γ) and K^+^ recycling cells (TAL-α) may account for the observation that 30% of NaCl reabsorption remains in mice with ROMK deleted (41).

Although there is clear mosaicism regarding cells that express ROMK at the apical membrane and those that don’t, all TAL cells exhibit message for *Kcnj1* (ROMK), as documented through both RNA-seq and in situ hybridization. This suggests that there is a pool of *Kcnj1* as a functional reservoir to adapt to altered reabsorptive work load or potassium homeostasis (42, 43). There is convincing evidence that diluting segments adjust K^+^ transport to meet homeostatic needs. In the amphiuma, dietary potassium adaptation increases the number of LBC cells (10), which according to our findings, express apical ROMK. Guggino and colleagues found that K^+^-loading changed the ratio of HBC to LBC cells (10) and suggested that K^+^-adaptation along this segment is mediated by cell type conversion rather than cell activation. Imai and colleagues reported that the ratio of LBC to HBC in hamster was greater in the cortical TAL, where net K^+^ secretion, rather than reabsorption, occurs (11). Finally, Sansom and colleagues reported that the loop of Henle of mice converts from a net potassium reabsorbing segment to one that secretes potassium when animals consume a low sodium/high potassium diet (44). As both TAL-β and TAL-γ cells lack ROMK at baseline, it will be interesting to determine whether both cell types contribute to potassium adaptation.

The present work extends recent transcriptomic analyses of human biopsies (45) and dissected mouse cortex (18), which typically identify two TAL clusters distinguished by the expression of Cldn10 and Cldn16. The nomenclature used for TAL clusters and their gene signatures, however, has not been consistent (17, 18, 20, 45). Our current results linking the K^+^ channel mosaicism with claudin expression, therefore allow us to propose a cell type classification that provides substantial explanatory power. For example, although early work indicated that CaSR activation causes a diuresis (46) by inhibiting ROMK (47), more recent studies using highly selective CaSR agonists and antagonists show that activation or inhibition of the CaSR along the TAL can alter Ca^2+^ reabsorption independent of changes in transepithelial voltage (6, 8). At least part of this effect involves regulation of claudin 14, which interacts with claudins 16 and 19 thereby reducing Ca^2+^ permeability (48). Our results show that genes for the CaSR, the parathyroid hormone receptor, and the vitamin D receptor are differentially expressed in TAL-β cells. The fact that we did not see Cldn14 likely reflects its limited expression in normocalcemic mice (49).

Our results further support the dominance of the circumferential Cldn16-positive tight junctional belt in TAL-β HBC cells, as they were arranged singly or in patches. While TAL-α cells expressed Cldn10 when they are located next to each other, Cldn16 was observed where TAL-α and TAL-β cells were adjacent (**Figure 3**). Dominance is reflected by the share of cortical Cldn16 in total tight junctional length which ranged between 37% and 97% in isolated tubules, suggesting its pronounced role in cTAL (15). The molecular cause for Cldn16 dominance remains unexplained, but it is also reflected in the widespread expression of Cldn16 at all cell borders when Cldn10 is deleted in mice (50) and in human HELIX syndrome, in which mutations in *CLDN10* lead to hypermagnesemia and nephrocalcinosis (35).

A small snRNA-seq cluster was identified here as well as in previous work (16, 17, 29, 45, 51) that expressed high levels of *Nos1,* identifying it as macula densa. Immunoreactivity for Cldn10 dominated in macula densa in agreement with earlier findings (24) and apical ROMK was mostly positive, but variable in strength as was another marker of macula densa, *Cox2* (52). Thus, along with Cldn16 absence, it may be concluded that the macula densa cell phenotype shares TAL-α properties (24).

The finding that pNKCC was present primarily in cells that lack ROMK, at least in rat, is surprising. Luminal insertion and activation of NKCC2 may be more challenged in ROMK-negative, than in ROMK-positive TAL cells chiefly of ISOM and OSOM; differences in the respective intracellular Cl^-^ concentration may be involved (53); failure to detect this in mouse and human TAL may be epitope-related (54).

Although it has been suggested that ROMK-positive cells correspond to microvilli-enriched or rough (R) as opposed to smooth (S) cells with few microvilli (11), we could not confirm that ROMK expression was linked to structurally distinct R or S cells. Although we could reproduce the heterogeneity in axial cell structure, we failed to confirm mosaic arrangement of S- and R-cells intra-segmentally, i.e., in any given region of interest within mTAL or cTAL (55).

**In conclusion**, we show that there are three distinct cell types in TAL (TAL-α, TAL-β, and TAL-γ) that are arranged in mosaic pattern across the TAL and conserved across species. Anchored in both the mosaicism of K^+^ channel expression and claudin expression, we assign independent regulation of paracellular routes for Na^+^ or Ca^2+^/Mg^2+^ to specific luminal or basolateral membrane proteins involved in transepithelial transport. These results show how one population of cells recycles K^+^ across the apical membrane, generates the transepithelial voltage, and may mediate K^+^ secretion, whereas another population, (TAL-β cells) with high basolateral K^+^ conductance and expressing Cldn16, transports Ca^2+^ and Mg^2+^. Interestingly, TAL-γ cells, only along the ISOM, have some characteristics of each of the other types and seem poised to mediate both transcellular NaCl and K^+^ reabsorption (**Figure 7)**. Diversity of TAL cell types, a feature conserved across species as diverse as amphiuma and human, therefore enables inter- and intra-segmental heterogeneity in TAL to provide homeostasis of mono- and divalent cation transport at steady state and when challenged.

## Methods

### Tissues

For animal study, adult male C57BL/6 mice (6 weeks old, n=5; Janvier labs, Le Genest-Saint-Isle, France) and male Long Evans rats (6 weeks old, n=5; Janvier labs) were used.

During housing, animals were kept in a 12-hour day/night cycle, with ad libitum access to drinking water and standard rodent chow (Altromin 1324). Animals were anesthetized with a mixture of ketamine (Pfizer, Karlsruhe, Germany) and xylazine (Rompun, Bayer, Germany). After opening the abdominal cavity, kidneys were perfusion-fixed via the abdominal aorta, first with 3% hydroxyethyl starch in 0.1 M Na-cacodylate (Caco) for 20 to 30 sec, then with 3% paraformaldehyde/3% hydroxyethyl starch in Caco for 5 min. Kidneys were removed and either incubated in 800 mosmol/kg H_2_O sucrose in PBS for 18 h, shock-frozen in liquid nitrogen-cooled isopentane and stored at −80°C, or postfixed in 3% PFA in PBS overnight, dehydrated via graded ethanol series to xylene, and embedded in paraffin. Samples were immersion-fixed in 3% PFA in PBS for 12 h and processed for paraffin embedding.

### Electron Microscopy

For ultrastructural analysis, kidney tissue was postfixed overnight at room temperature in 1.5% glutaraldehyde/1.5% paraformaldehyde containing 0.05% picric acid in Caco, then in 1% osmium tetroxide/0.8% potassium hexacyanoferrate in Caco for 1.5 h at room temperature (RT) for TEM or in 1% aqueous osmium tetroxide for SEM. Tissues were then dehydrated and embedded in epoxy resin for semithin sectioning and light microscopy (LM) or ultrathin sectioning and TEM analysis using standard methodology. For SEM, samples were high pressure-critical point-dried and sputter-coated. For pre-embedding immunoperoxidase staining 30 μm-thick cryostat sections were cut from frozen kidneys, stained with anti-ROMK antibody and prepared for electron microscopic analysis as previously described (52).

### Immunofluorescence

Cryostat kidney sections (6 µm thickness) were cut using a CM 3050S microtome (Leica, Wetzlar, Germany) and permeabilized for 30 min with 0.5% Triton X-100 (Merck, Darmstadt, Germany) in PBS. Paraffin sections (4 µm thickness) were prepared with an RM 2125 RT microtome (Leica), deparaffinized in xylene, and rehydrated via graded ethanol series. Heat-induced epitope retrieval was performed for 6 min in citrate buffer (pH 6.0) using a pressure cooker. Before antibody incubation, cryosections and paraffin sections were blocked with 5% skim milk (Difco, BD, USA) for 30 min. Primary antibodies were then applied overnight at 4°C (**Supplemental Table 2**). Secondary antibodies were applied for 60 min at room temperature. Nuclei were counterlabeled with DAPI (Sigma-Aldrich). Sections were mounted in 1:9 PBS:glycerol or ProLong Glass Antifade Mountant® (ThermoFisher Scientific) and examined using LSM 5 Exciter (Carl Zeiss, Jena, Germany) or SP8 (Leica) confocal microscopes. The percentages of ROMK-, Kir4.1-, pNKCC2-negative cells in TAL were quantified in transversely cut tubules and expressed as percentages of the total numbers of TAL cells counted per kidney zone. At least 20 similar tubular profiles were evaluated in each zone (inner stripe, outer stripe, cortex).

### In situ hybridization

*In situ* hybridization was performed on perfusion-fixed, paraffin-embedded rat kidney tissue using an RNAscope® 2.5 HD Brown Reagent Kit (Advanced Cell Diagnostics ACD, Hayward, CA) according to manufacturer’s instructions. Briefly, dewaxed paraffin sections (4 µm thickness) were incubated with peroxidase blocker and boiled for 30 min at 100°C in a pre-treatment solution. Sections were treated with protease for 30 min at 40°C. A ROMK target probe (ACD) was hybridized at 40°C for 2 h in the dark using the HybEZ Hybridization System (ACD). After several amplifying and washing steps, the sections were stained with chromogenic substrate for detection of hybridization signals. ROMK mRNA signal was identified as red punctate dots. Nuclei were counterstained with hematoxylin for 2 min at RT. Sections were coverslipped with DakoCytomation Glycergel mounting medium. Sections were examined with a Leica DMRB microscope equipped with an AxioCam MRc camera (Zeiss). Endogenous housekeeping gene *UBC* was used as a positive control; bacterial *dapB* served as a negative control to assess background signal.

### Electrophysiology

C57BL/6J mice (8 weeks old) were purchased from the Jackson Laboratory (Bar Harbor, ME). Mice were euthanized by CO_2_ inhalation plus cervical dislocation. Their abdomen was opened to expose the left kidney. Kidney was perfused with 2 ml L-15 medium (Life Technology) containing Type 2 collagenase (250 units/ml), then removed and the cortex dissected into small pieces for additional incubation in collagenase-containing L-15 media for 30-40 min at 37°C. Pieces were then washed three times with fresh L-15 medium and transferred to an ice-cold chamber for dissection. cTALs were isolated as described (21). Isolated cTALs were placed on a cover glass coated with polylysine. The cover glass was then transferred into a chamber mounted on an inverted microscope.

### Patch-clamp Experiments

We used an Axon 200A amplifier to conduct the whole-cell voltage clamp experiments in isolated and split-open cTAL. The pipette solution contains 140 mM KCl, 2 mM MgATP, 1 mM EGTA, and 10 mM HEPES (pH 7.4) for measurement of K^+^ currents. The bath solution was similar to the pipette solution without MgATPThe currents were low-pass filtered at 1 kHz, digitized by an Axon interface with a 4-kHz sampling rate (Digidata 1440A). Data were analyzed using the pClamp software system 9.0 (Axon).

### Membrane Voltage Measurement

C57 Bl/6 were euthanized, the kidneys removed and sliced for enzymatic tubule preparation as described (56). Tubule suspension was obtained by dissociation of slices in a thermomixer (Eppendorf, 850 rpm, 37°C). The incubation solution was composed of (mM) 140 NaCl, 0.4 KH2PO4, 1.6 K2HPO4, 1 MgSO4, 10 Na-acetate, 1 α-ketoglutarate, 1.3 calcium gluconate, pH 7.4, and contained (mg/l) 25 DNase I, 375 glycine, 48 trypsin inhibitor and 2000 collagenase II. For TAL sorting, the incubation solution was supplemented with 500 mg/l albumin. TAL were identified, isolated at 4°C according to morphological characteristics (56), and used for single tubule immunofluorescence or membrane voltage measurements.

For immunofluorescence, single cTAL segments were placed on microscope slides (EprediaTM Polysine adhesion slides, Thermo Fisher Scientific) and fixed with 4% paraformaldehyde for 7 min, washed several times with 10 mM sodium citrate pH 6 and heated in this buffer for 3h at 70°C. After washing with PBS-T (0.3% Triton X-100 in phosphate-buffered saline) tubules were incubated with primary antibodies (**Supplemental Table 2**) in PBS-T + 5% BSA overnight at 4°C. Secondary antibody (**Supplemental Table 2**) was incubated for 1-2h after extensive washing with PBS-T. Confocal Airyscan images were acquired using Zeiss LSM 880 confocal laser-scanning microscope (excitation laser: 488, 594 and 633 nm) equipped with an Airyscan unit. Enzymatically dissected TAL were transferred into the bath chamber on the stage of a Leica Stellaris 5 confocal microscope (Leica microsystems, Wetzlar, Germany) at 37°C and perfused by a concentric glass pipette perfusion system. Perfused TAL were stained by basolateral as well as luminal incubation with the membrane voltage dye Di-8-ANEPPS (6 µM; D3167, Thermo Fisher Scientific) in the presence of 0.02% Pluronic F-127 for 30 min in control solution containing (mM) 145 NaCl, 0.4 KH2PO4, 1.6 K2HPO4, 1 MgCl2, 1.3 calcium gluconate, 5 glucose, pH 7.4). After staining, fluorescence (ex: 486 nm, em: 505-700 nm) was recorded every 5 seconds at symmetric control solution. Then the tubule was superfused with 30 mmol/L K^+^ solution containing (mM) 120 NaCl, 26.4 KCl, 0.4 KH2PO4, 1.6 K2HPO4, 1 MgCl2, 1.3 calcium gluconate, 5 glucose, pH 7.4) for 3 min. Fluorescence measurements were analyzed using the LasX Office software (Leica microsystems, Wetzlar, Germany) after background subtraction. 2-3 cells without movement artifacts were selected and fluorescence intensity was normalized to the mean intensity of 4 measurements preceding solution change. TAL cells were categorized into 2 groups, one with no increase in mean intensity in the presence of 30 K^+^ and another with mean intensity increase in the presence of 30 K^+^ upon 4 consecutive measurements.

### Analysis of TAL snRNA-seq Dataset

#### NKCC2-INTACT Mouse Model

Sodium-potassium-chloride cotransporter 2 (NKCC2)-Cre-INTACT mice were used for single-nucleus RNA sequencing (snRNA-Seq) experiments. The Cre/LoxP system was employed to label TAL nuclei with nuclear green fluorescent protein (GFP), enabling fluorescence-activated nuclei sorting (FANS) enrichment. This approach, previously described for the DCT (26), uses the INTACT system to fluorescently label the nuclei of genetically targetable cell populations. The INTACT system tags the C-terminus of SUN1, a nuclear membrane protein, with two tandem copies of superfolder GFP (sfGFP) and six copies of the Myc epitope (CAG-SUN1-sfGFP-Myc). To target TAL cells specifically, INTACT mice were crossed with NKCC2-CreERT2 mice (generated by Ruisheng Liu), producing a mouse line that expresses sfGFP at the inner nuclear membrane in TAL cells upon tamoxifen induction.

To induce sfGFP expression, NKCC2-Cre-INTACT mice were intraperitoneally injected with 1 mg tamoxifen (dissolved in corn oil) daily for 5 days, followed by a 10-day induction period. Mice were housed under standard conditions with a 12-hour light/dark cycle and food and water ad libitum. After tamoxifen induction, mice were anesthetized using isoflurane, followed by cardiac perfusion with 10–15 ml of phosphate-buffered saline (PBS) to remove blood cells. Kidneys were harvested, and the kidney capsules were removed with the ureters trimmed prior to dissection. Prior to sequencing, kidneys were manually dissected into cortical and medullary regions. Kidneys were sliced along the cortical-medullary axis in a Petri dish. A superficial strip (∼20% depth) from each slice was separated as the cortex zone, while the remaining tissue was collected as the medulla zone. The dissected cortex and medulla portions were snap-frozen in liquid nitrogen immediately after harvest and stored at −80 °C until further downstream nuclei isolation, fluorescence-activated nuclei sorting (FANS), and single-nucleus RNA sequencing (snRNA-Seq).

Nuclei were isolated using the 10x Genomics kit (Chromium Nuclei Isolation Kit PN-1000494) according to the manufacturer’s instructions. The nuclei suspension was mixed with 5µl Vybrant Ruby stain and incubated on ice for 15 min. The nuclei were sorted in 500 µl final resuspension buffer (FBS, 1ml DPBS with 1% BSA + 5 µl Protector RNase inhibitor) with a low flow rate and pressure to ensure high viability. Two main gates were used. A 561+683 emission for the ruby stain and a 488 530/40 emission for GFP. Low trigger pulse width was used as singlet discriminator. One hundred fifty thousand nuclei were collected and centrifuged at 500g for 5 min at 4°C. The top supernatant was carefully removed, and 6 µl volume was left to resuspend the nuclei pellet.

Single nucleus RNA-seq was performed using a PIPseq™ T10 3ʹ Single Cell RNA Kit v5.0 (Fluent BioSciences). Single nucleus suspensions with a maximal concentration of 3,400 nuclei/uL (for a total of 17,000 nuclei per sample) were added directly into Particle-templated Instant Partition (PIP) tubes. Following the manufacturers protocol, with subsequent vortexing the nuclei were partitioned into PIPs. Following capture and lysis of the RNA and downstream cDNA generation and amplification, the resulting cDNA was run on a 2% gel electrophoresis for approximate band size (greater than 500bp). The cDNA was then used as a template for library preparation according to the reagent kit protocol. https://www.fluentbio.com/resource-category/user-guides/

The dataset was processed using the Seurat pipeline, incorporating doublet removal with DoubletFinder, ambient RNA correction with SoupX, normalization via scTransform, and integration within Seurat. A 2D UMAP projection was generated using the first 10 principal components to visualize cellular clusters (27). Cortex and Medulla scores were created using Seurat’s AddModuleScore function, based on the average expression of the top genes within cortex and medulla. Dimension, feature, dot, and violin plots were generated using Seurat’s built-in visualization functions.

The final enriched-TAL dataset included 31,884 features across 11,724 samples, comprising 4,076 TAL cortical nuclei with a median of 1,366 genes and a mean of 2,162 reads per nucleus, and 7,648 TAL medullary nuclei with a median of 1,412 genes and a mean of 2,320 reads per nucleus.

The previously published mouse dataset was downloaded from GEO (GSE129798) (16). Sample information was extracted from the data embedded into the cell barcodes. This dataset utilized whole-kidney single-cell suspensions from the cortex (Zone 1), outer medulla (Zone 2), and inner medulla (Zone 3) of two adult (8–9 weeks old) male and two adult female C57BL6/J mice. The human dataset from the KPMP was downloaded from 13 healthy references on November 29, 2022 (57). This dataset utilized single-nuclei suspensions from protocol renal biopsies, percutaneous nephrolithotomy biopsies, and living donor nephrectomy tissues. The samples contained spatial information, which KPMP determined by analyzing 5 µm histology sections adjacent to the sampled areas, reporting the relative composition of the cTAL and mTAL. The rat dataset was downloaded from GEO (GSE209821) (51). This dataset utilized whole-kidney single-nuclei suspensions from three male lean ZSF1 rats.

The whole-kidney datasets were analyzed using the Seurat Standard Workflow (Seurat v4.3.0, (58). For the mouse and rat datasets, the TAL cluster was identified by the abundant and specific expression of the TAL genetic marker *Slc12a1*. In the human dataset, the TAL was identified by subsetting cTAL, mTAL and MD cells, following the original KPMP nomenclature. Erroneous cell clusters were removed prior to final reprocessing through the Seurat Standard Workflow. Profiles of Slc12a1-positive cells were analyzed using the Seurat R package through unsupervised clustering. Dimensionality reduction analysis was performed via Principal Component Analysis (PCA). Clusters were generated employing Seurat’s built-in commands, FindNeighbors and FindClusters. The data were visualized as a two-dimensional Uniform Manifold Approximation and Projection (UMAP) plot. Clusters were annotated based on prominent *Cldn10*, *Cldn16*, and *Nos1* gene expression, as identified using the FindMarkers function. As a result, the clusters were named TAL-α, TAL-β, and MD cell types. The final Seurat object for the mouse, derived from four animals, comprised a total of 31,884 features across 1,529 samples. The final Seurat object for humans, derived from thirteen subjects, comprised a total of 53,884 features across 9,410 samples. The final Seurat object for the rat, derived from three animals, comprised a total of 39,161 features across 8,408 samples. Differential gene expression analysis was conducted using Seurat’s FindMarkers function, ordering the resulting dataframe according to the log2FC and an adjusted p-value of 0.05. Plots were generated using Seurat’s built-in visualization functions.

#### Sex as a Biological Variable

Both sexes were employed for these studies. The results describe those findings shared by sexes and those where there are differences.

#### Statistics

GraphPad Prism8 software was used to analyze parameters. Quantification of cell counting data is presented as the mean ± standard deviation. Evaluation of multiple groups was performed using ANOVA followed by Tukey’s post hoc test A significance level of P < 0.05 was accepted as significant.

#### Study Approval

Animal studies in Germany were approved by the Berlin Senate (animal experimental authorization G0285/10). Animal studies in Oregon were approved by the Oregon Health and Science University IACUC (protocol IP00286). Animal studies in New York were approved by New York Medical College (IACUC#21127). Human kidney samples were obtained from morphologically normal portions of tumor nephrectomy specimens after written consent (Charite No. EA4/002/24).

## Supporting information

Supplemental DEGs

## Data, Materials, and Software Availability

PIPseeker output files and an annotated Seurat Object are uploaded to GEO (GSE284450; reviewer token: cbypsgqylxknhyj). To ensure data accessibility we also made the TAL-INTACT dataset available for further exploration via an interactive web tool generated using ShinyCell^17^ at https://ellisonlab.shinyapps.io/tal_shinycell/ Previously published data were used for this work: References (51), (16), (29). All study data are included in the article and/or SI Appendix. Code for data processing and figure generation is available on GitHub (https://github.com/OHSU-NHT).

## Author contributions

H.D, J.B.L, D.H.E., and S.B. designed research; H.D., J.B.L, A.S., X.T.S., J.N., C.L.Y, J.C., X.P.D, Y.S., D.E.Y, C.Q., N.H., M.B., and S.B. performed research; H.D., J.B.L, A.S., X.T.S., J.N., C.L.Y, J.C., X.P.D, Y.S., D.E.Y, C.Q., N.H., M.B., and S.B. analyzed data; K.E. and B.E. developed the PIP Seq model; R.L. generated the NKCC2-CRE mouse model; H.D, J.B.L, D.H.E., and S.B. wrote the paper.

## Acknowledgements

This work was financially supported by Deutsche Forschungsgemeinschaft BA700/22-2, MU2924/2-2, and SFB 1365-C04/-S01 to SB. It was also supported by the National Institutes of Health R01 DK133220, DK51496, U54TR001628 and a Fondation LeDucq: Transatlantic Network of Excellence 17CVD05. HD was supported by a doctoral fellowship from Ministry of National Education, Turkey. JBL was supported by a Benjamin J. Lipps Research Fellowship from KidneyCure. JNC is supported by NIH T32HL094294.JWN is supported by NIDDK DK121737. We thank Kerim Mutig for scientific advice, Anette Drobbe for secretarial help, Kerstin Riskowsky for expert technical help and Sara Timm, Petra Schrade, and John Horn (Core Facility for EM) for help with microscopy. We are grateful to Katharina Broeker for methodological help in RNAscope technology.

