## Supplemental figures for "Distinct cell types along thick ascending limb express pathways for monovalent and divalent cation transport"

**Supplemental Information Appendix**

**Supplemental Table 1: Differentially expressed genes list**

**Supplemental Table 2. Primary and secondary antibody list**

| **Primary and secondary antibodies** | **Host & Clonality** | **Company** | **Dilution for IHC** |
|---|---|---|---|
| Anti-CaSR | Rabbit, polyclonal | Enzo, Life Science | 1:500 |
| Anti-claudin 10 | Rabbit, polyclonal | Invitrogen | 1:200 |
| Anti-claudin 16 | Rabbit, polyclonal | BiCell Scientific | 1:100 |
| Anti-claudin 16 | Mouse, monoclonal | Provided by H. Dimke | 1:100 |
| Anti-claudin 16 | Rabbit, polyclonal | J.H. laboratory, Washington University, St. Louis | 1:100 |
| Anti-Cox-2 | Goat, polyclonal | Santa Cruz Biotechnology | 1.300 |
| Anti-Kir4.1 (KCNJ10) | Rabbit, polyclonal | Alomone Labs | 1:500 |
| Anti-NKCC2 | Guinea pig, polyclonal | Provided by David H. Ellison | 1:5000 |
| Anti-nNOS | Rabbit, polyclonal | Thermo Fisher Scientific | 1.1000 |
| Anti-phospho-NKCC2 (Ser 130) | Sheep, polyclonal | MRC Dundee | 1:2000 |
| Anti-phospho-NKCC2 (Thr 96) | Rabbit, polyclonal | Provided by K. Mutig | 1:1000 |
| Anti-ROMK (KCNJ1) | Rabbit, polyclonal | Alomone Labs | 1:500 |
| Cy3-coupled donkey anti-rabbit IgG | Donkey, polyclonal | Dianova | 1:250 |
| Cy2-coupled donkey anti-rabbit IgG | Donkey, polyclonal | Dianova | 1:100 |

| | | | |
|---|---|---|---|
| Cy3-coupled donkey anti-mouse IgG | Donkey, polyclonal | Dianova | 1:250 |
| Cy2-coupled donkey anti-mouse IgG | Donkey, polyclonal | Dianova | 1:100 |
| Cy3-coupled donkey anti-guinea pig IgG | Donkey, polyclonal | Dianova | 1:250 |
| Cy2-coupled donkey anti-guinea pig IgG | Donkey, polyclonal | Dianova | 1:100 |
| Cy3-coupled donkey anti-sheep IgG | Donkey, polyclonal | Dianova | 1:250 |
| Cy2-coupled donkey anti-sheep IgG | Donkey, polyclonal | Dianova | 1:100 |
| Donkey anti-rabbit IgG (H+L) Alexa 488-conjugated | Donkey, polyclonal | Invitrogen | 1:400 |
| Goat-anti-mouse Alexa-488 | Goat, polyclonal | Thermo Fisher Scientific | 1:250 |
| Goat-anti-guinea pig Alexa-594 | Goat, polyclonal | Thermo Fisher Scientific | 1:250 |
| Goat-anti-rabbit Alexa-633 | Goat, polyclonal | Thermo Fisher Scientific | 1:250 |
| Anti-rabbit immunoglobulins HRP | Swine, polyclonal | Dako | 1:2000 |
| Affinity pure Fab fragment goat anti-rabbit IgG | Goat, polyclonal | Jackson ImmuneResearch Laboratories | 1:20 |

**Supplemental Figure 1. ROMK immunostaining in TAL across species and gender; distribution of pNKCC2, rat kidney. (A)** ROMK immunostaining in rat, mouse and human kidney samples is identified by mosaic luminal signal in TAL double-stained for continous NKCC2 signal; DAPI blue nuclear staining in merge images. DIC, differential interference contrast optics. Bars in epithelia identify cell borders between ROMK-positive and ROMK-negative cells. **(B-D)** Numerical evaluation of ROMK-negative cells in TAL of male **(B)** and female **(C)** mice and female rats **(D)**; values are means ± SD from n=5 mice or rats. * $P < 0.05$; ** $P < 0.01$; *** $P < 0.001$; **** $P < 0.0001$ by ANOVA and post-hoc comparisons. **(E)** ROMK and phospho-(p)-Ser130-NKCC2 signals reveal mutually exclusive mosaicism across renal zones by double immunostaining. **(F)** Numerical evaluation of pNKCC2-negative cells cells across the zones. **** $P < 0.0001$; ns, not significant ns, not significant by ANOVA and post-hoc comparisons.

**Supplemental Figure 2. ROMK mRNA expression in mTAL, rat kidney. (A,B)** ROMK mRNA is visualized by in situ hybridization using RNAscope technology. All cells are labeled in TAL profiles (asterisks) and collecting ducts; no staining in intercalated cells is indicated by arrowheads **(A)**. No signal is obtained from a control bacterial DapB probe **(B)**. Scale bars indicate magnification.

**Supplemental Figure 3. Macula densa-specific expression patterns, rat and mouse kidney**. **(A)** Rat macula densa staining by (nicotinamide adenine dinucleotide phosphate) diaphorase associates with ROMK and NKCC2 immunorective signals. **(B)** ROMK and cyclooxygenase-2 (COX-2) immunoreactive signals are colocalized in macula densa. **(C)** ROMK and neuronal nitric synthase (nNOS) immunoreactive signals are colocalized in macula densa; note weak ROMK signal occurring in places. **(D)** In mouse kidney, macula densa ROMK immunoreactive signal may be weak or displaying scattered positivity, here in double staining for nNOS. Scale bars indicate magnification.

**Supplemental Figure 4. Distribution of Claudins 10 and 16 and Kir4.1 in isolated mouse cTAL.** Triple staining immunohistochemistry shows one isolated TAL segment; note that Claudin (Cldn)10 staining is mutually exclusive to Cldn16 and basolateral Kir4.1 staining. Scale bar indicates magnification.

**Supplemental Figure 5. Cell type distribution and kidney zone localization of mouse Thick Ascending Limb (TAL) cells as determined by single-nucleus RNA sequencing.** **(A)** Uniform Manifold Approximation and Projection (UMAP) plot showing Slc12a1 transcript levels, a canonical marker for TAL cells. **(B)** UMAP plot indicating the anatomical origin of TAL cells, classified into manually dissected kidney zones: cortex and medulla. **(C)** UMAP plot identifying distinct TAL cell populations. TAL-α accounts for 55.8%, TAL-β for 18.4%, TAL-γ for 18.8%, macula densa (MD) cells for 6.86%, and proliferative (Prolif) cells for 0.14%. **(D)** Heatmap illustrating the expression of canonical markers across all kidney cell types. **(E)** Bar graph displaying the percentage distribution of TAL subtypes across kidney zones. In the cortex: TAL-α comprises 41.4%, TAL-β 34.4%, TAL-γ 0.6%, MD 17.4%, and proliferative cells 0.2%. In the medulla: TAL-α represents 60.3%, TAL-β 9.9%, TAL-γ 28.5%, MD 1.2%, and proliferative cells 0.1%. **(F)** UMAP plot showing the distribution of Kcnt1 expression across kidney zones. **(G)** Dot plot of the top 10 differentially expressed genes (DEGs) in each TAL cell population. Gene expression was normalized and z-score scaled to compare relative expression across clusters. "Average expression (Avg Expr)" refers to the z-scored of the average gene expression within a cluster, while "Percent expressed (Pct Expr)" denotes the proportion of cells within a cluster expressing the gene.

**Supplemental Figure 6. Highlighted gene expression profiles from enriched single-nucleus RNA sequencing analysis.**

Dot plots displaying the expression of selected gene categories in each TAL cell population: (**A**) Solute carriers, (**B**) Kinases, (**C**) Phosphatases, (**D**) Phosphodiesterases (PDEs) and ubiquitin ligases, (**E**) Genes associated with ionic transport and regulation, (**F**) Cellular receptors and signaling pathways, and (**G**) Nuclear receptors and transcription factors. Gene expression was normalized and z-score scaled to facilitate comparisons across clusters. "Average expression (Avg Expr)" refers to the z-scored of the average gene expression within a cluster, while "Percent expressed (Pct Expr)" denotes the proportion of cells within a cluster expressing the gene.

**Supplemental Figure 7. Cortico-Medullary TAL markers from enriched Single-Nuclei RNA-Seq analysis.**

(**A, B**) UMAP projections showing the cortex (**A**) and medulla (**B**) score expression across all nuclei. Cortex and medulla scores were created using Seurat's ‘AddModuleScore’ function, based on the average expression of top differentially expressed genes (DEGs) enriched in the cortex ((*Pde10a*, *Sfrp1*, *Car15*, *Enox1*, *Pth1r*, *Cadm2*, *Thsd4*, *9530026P05Rik*, *Cdh6*, *Sec14l1*, *Klk1*, *Tmem117*, *Clcnkb*) and the medulla (*Ranbp3l*, *Sgcz*, *Enpp2*, *Pcsk6*, *Magi2*, *2210408F21Rik*, *Gm16310*, *Mrps6*, *Cacnb4*, *Slc5a3*, *Pax2*, *Apbb2*, *Scin*). Color intensity represents the relative expression of these scores across nuclei. The spatial arrangement of the clusters reflects a continuous anatomical progression consistent with the organization of the kidney, highlighting the ability of these scores to distinguish cortical and medullary cells. This approach provides a valuable tool for analyzing TAL cells in the absence of spatial data. (**C, D**) Violin plots showing the expression of the top DEGs used to create cortex (**C**) and medulla (**D**) scores.

**Supplemental Figure 8. Cell type distribution and kidney zone localization of mouse TAL cells as determined in previously published Whole-Kidney single-cell RNA-seq Dataset by Ransick (1)**

TAL cells (*Slc12a1* +) were subsetted from a previously published Whole-Kidney single-cell

RNA-seq dataset by **(1)** (**A**) Uniform manifold approximation and projection (UMAP) displaying the origin of cells from different kidney zones according to **(1).** Z1 corresponds to the cortex, Z2 corresponds to the outer medulla, and Z3 corresponds to the inner medulla. Graph also shows percentage of cell type in different kidney zones. Z1, TAL-α (50.8%), TAL-(42.6%), MD (6.6%); Z2, TAL-α (77%), TAL-β (22.6%), MD (0.4%); Z3, TAL-α (98.9%), TAL-β (1.1%). (**B)** Dot plot of the top differentially expressed genes (DEGs) in each TAL cell population. UMAP plot displaying *Kcnj10* (**C**), *Cldn10* (**D**), *Cldn16* (**E**) transcript levels. Dot plot illustrating the distributions of transcripts associated with sodium transport (**F**), calcium & magnesium (**G**) and potassium (**H**) transport**.** Genes related to the regulation of sodium transport exhibit higher expression in TAL-α, while genes linked to calcium transport show higher expression in TAL-β. (**I**) Dot plot illustrating the distributions of *Avpr2*, *Pth1r*, *Clcnkb* and *Clcnka* transcripts levels across kidney zones. *Avpr2* exhibit higher expression in TAL-α with peak levels in medullary regions, while *Pth1r* show higher expression in TAL-β with peak levels in cortex. *Clcnka* and *Clcnkb* serve as cortico-medullary markers, with *Clcnkb* more express in cortical regions and *Clcnka* in medullary regions (2). The data is normalized and scaled using a z-score to compare relative expression levels among cell clusters. "Average expression (Avg Expr)" refers to the z-scored of the average gene expression within a cluster, while "Percent expressed (Pct Expr)" denotes the proportion of cells within a cluster expressing the gene. (**J**) Graph showing percentage of cell types in males and females. Females, TAL-α (68.8%), TAL-β (29.6%), MD (1.6%); Males, TAL-α (82.6%), TAL-β (16.4%), MD (1.0%).

**Supplemental Figure 9. Cell type distribution and kidney zone localization of human TAL cells as determined in previously published single-nuclei RNA-seq Dataset by (3)** (**A**) Uniform manifold approximation and projection (UMAP) visualization of TAL cell populations identified from a single-nucleus RNA-sequencing (snRNA-seq) dataset (WashU-

UCSD_HuBMAP_KPMP-Biopsy_10X-R_12032021.h5Seurat) obtained from the Kidney Precision Medicine Project (KPMP) (3). The TAL population consists of three distinct cell types, all characterized by *SLC12A1* expression: TAL-α, TAL-β, and macula densa (MD). TAL-α accounts for 73.9%, TAL-β for 17.9% and macula densa (MD) cells for 8.2%. (**B-C**) UMAP projections displaying nuclei expressing the genes claudin 10 (*CLDN10*) (**B**) and claudin 16 (*CLDN16*) (**C**). (**D**) UMAP projection illustrating the zonal origin of TAL cells across different kidney regions, as determined from KPMP. (**E-G**) Dot plots and heatmaps illustrate gene expression patterns associated with: (**E**) Sodium transport (*CLDN10, WNK4, WNK1, STK39, PTGER3, SLCO3A1, AVPR2*), which are enriched in TAL-α. (**F**) Calcium and magnesium transport (*C LDN16, CLDN19, CLDN14, CASR, PTH1R, VDR, CNNM2)*, with higher expression in TAL-β. (**G**) Potassium transport (KCNMA1, KCNJ1, KCNJ10, KCNJ16, KCNT1). Data normalization and scaling: Gene expression data were normalized and z-score scaled to enable comparison of relative expression levels across clusters. "Avg Exp" (Average Expression) and “Expression” (Relative Expression) represents the z-scored mean gene expression within a cluster, while "Pct Exp" (Percent Expressed) denotes the percentage of cells in a cluster with detectable expression of the gene. (**H**) Volcano plot showing differentially expressed genes (DEGs) that define the TAL-α and TAL-β cell clusters. (**I**) Correlation analysis comparing the cluster-defining DEGs of TAL-α and TAL-β between human (3) and mouse (1) datasets. The x-axis represents the average log2 fold change of human DEGs, while the y-axis represents the average log2 fold change of mouse DEGs. The coefficient of determination ($R^2$) is provided. (**J**) Graph showing percentage of cell types in males and females. Females, TAL-α (75.1%), TAL-β (19.6%), MD (5.3%); Males, TAL-α (73.1%), TAL-β (16.8%), MD (10.1%).

**Supplemental Figure 10. Cell type distribution of rat TAL cells as determined in previously published Whole-Kidney single-nuclei RNA-seq Dataset by (4).**

TAL cells (*Slc12a1* +) were subsetted from a previously published Whole-Kidney single-cell RNA-seq dataset by **(4).** Uniform manifold approximation and projection (UMAP) projection displaying nuclei expressing *Slc12a1* **(A),** *Cldn10* (**B**), *Cldn16* (**C**) and *Nos1* (**D**) genes from single-nuclei RNA-sequencing (snRNA-seq) rat dataset **(4)**. (**E**) DotPlot of genes 'more expressed in the cortex', *Enox1*, *Clcnkb*, and genes 'more expressed in the medulla', *Clcnka*, *Slc4a7*, *Ank2*, *Cryab*. (**F**) UMAP projection of thick ascending limb (TAL) population cells. There are four clusters among *Slc12a1* positive nuclei, TAL-α1, TAL-α2, TAL-β, and macula densa (MD) cell types**.** TAL-α accounts for 83.7 % (comprising TAL-α1 (32.8%) and TAL-α2 (50.9%)), TAL-β for 14% and macula densa (MD) cells for 2.3%. **(G-I)** Dot plot illustrating the distributions of transcripts associated with sodium transport (**G**), calcium transport (**H**) and potassium transport (**I**). Genes related to the regulation of sodium transport exhibit higher expression in TAL-α, while genes linked to calcium transport show higher expression in TAL-β. The data is normalized and scaled using a z-score to compare relative expression levels among cell clusters. "Average expression (Avg Expr)" refers to the z-scored of the average gene expression within a cluster, while "Percent expressed (Pct Expr)" denotes the proportion of cells within a cluster expressing the gene. (**J**) Volcano Plot showing cluster-defining differentially expressed genes (DEGs) of rat TAL-α (TAL-α1 and TAL-α2) and TAL-β cell types. (**K**) Correlation analysis of cluster-defining DEGs between rat (4) and mouse (1). TAL-α versus TAL-β. The x axis displays the average log2 fold change of rat DEGs, while the y axis shows the average log2 fold change of mouse DEGs. R2 represents the coefficient of determination.

**Supplemental Figure 11. Anatomically stratified TAL cell type characterization from transcriptomic analysis and expression of key membrane proteins.** Data are from single-cell RNA-sequencing (scRNA-seq) and immunohistochemistry (IHC). Renal parenchymal zones are distinctly shaded. On the right, cells labeled red are positive for the respective product, cells labeled white are negative. Tables below list the key proteins and mRNAs in assignment to the respective cell types and kidney zones.

**Supplemental Figure 1. ROMK immunostaining in TAL across species and gender; distribution of pNKCC2, rat kidney**

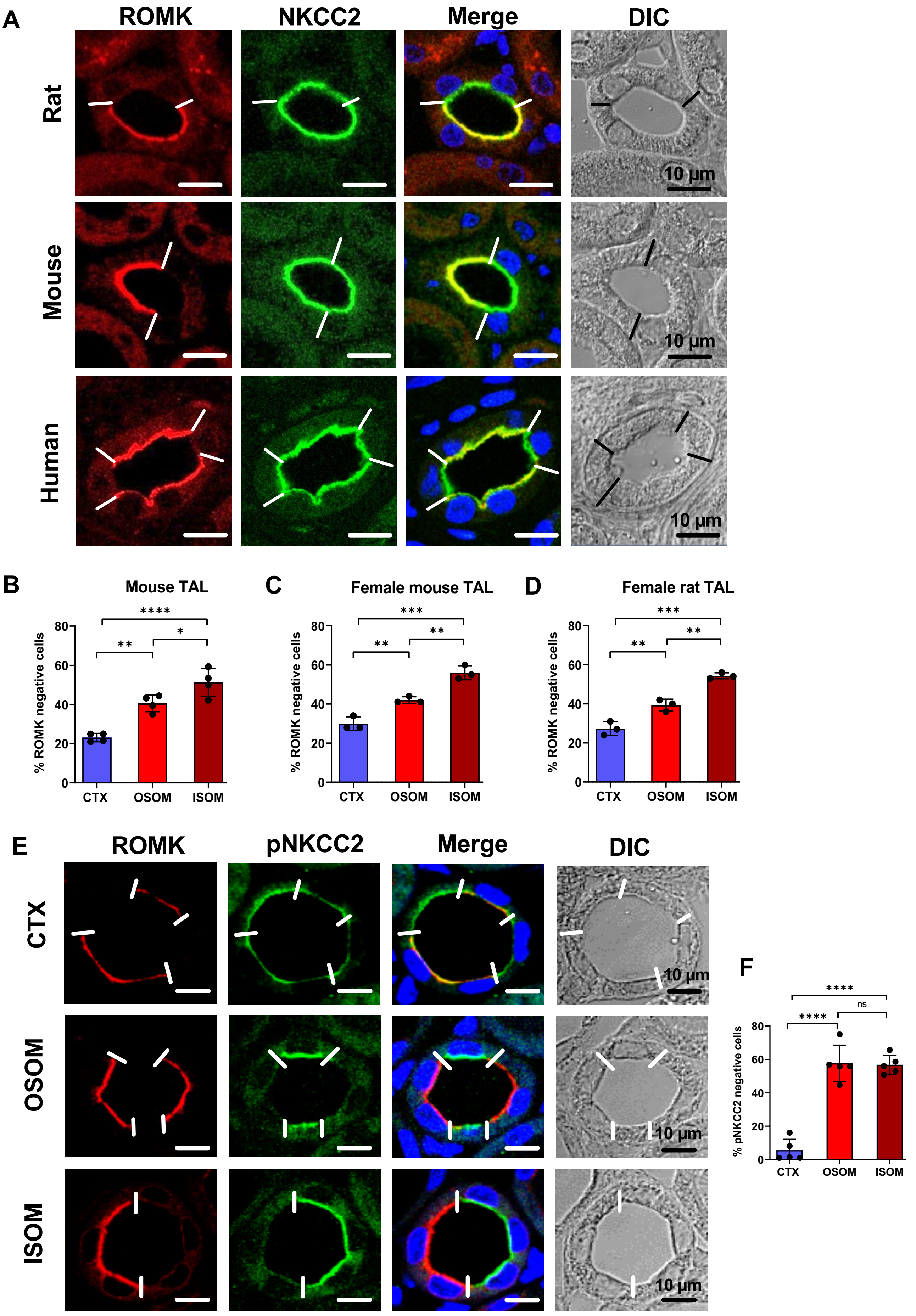

**Supplemental Figure 2. ROMK mRNA expression in mTAL, rat kidney**

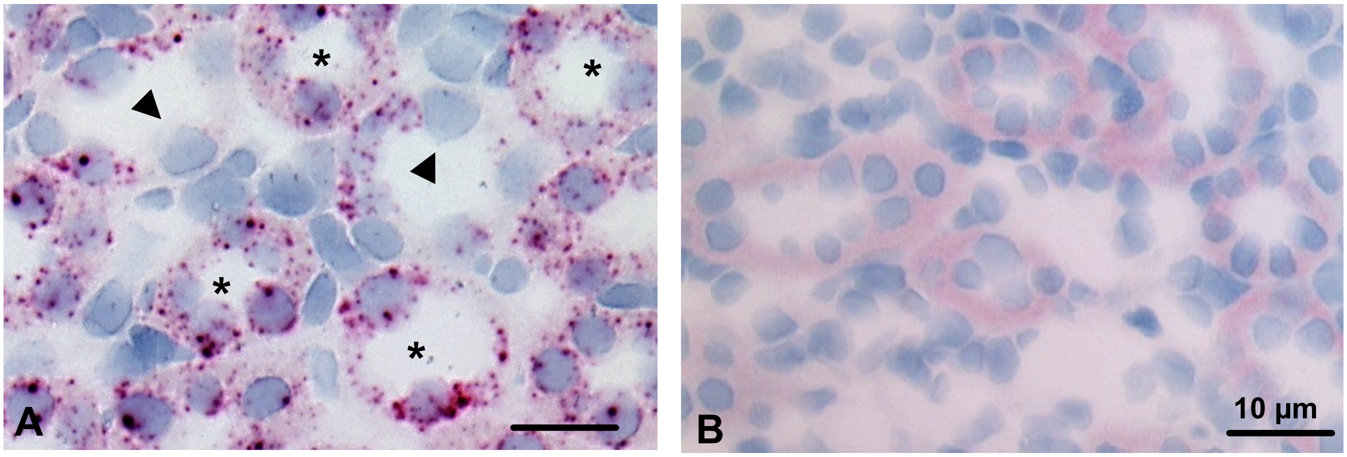

**Supplemental Figure 3. Macula densa-specific expression patterns, rat and mouse kidney**

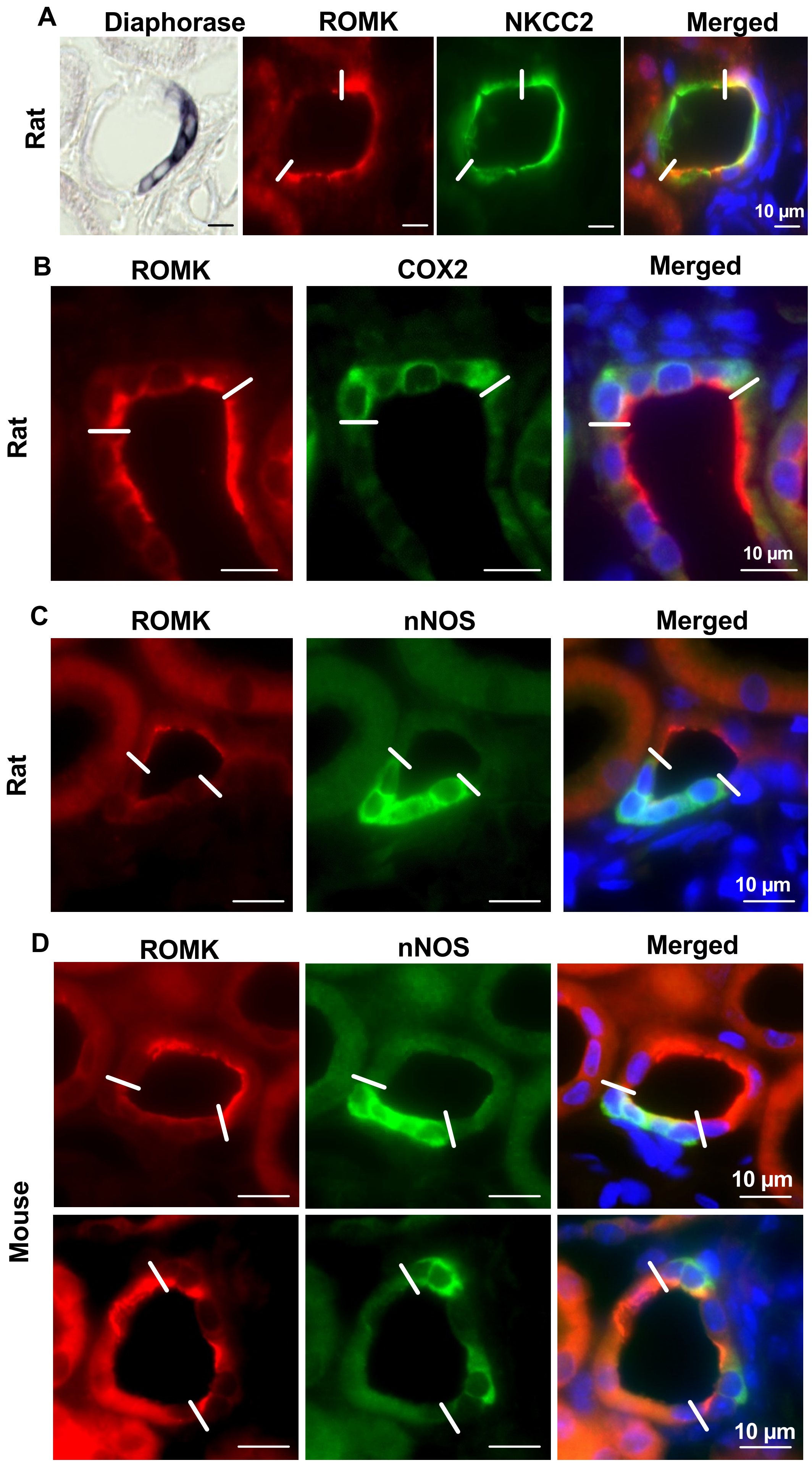

**Supplemental Figure 4. Distribution of Claudins 10 and 16 and Kir4.1 in isolated mouse cTAL**

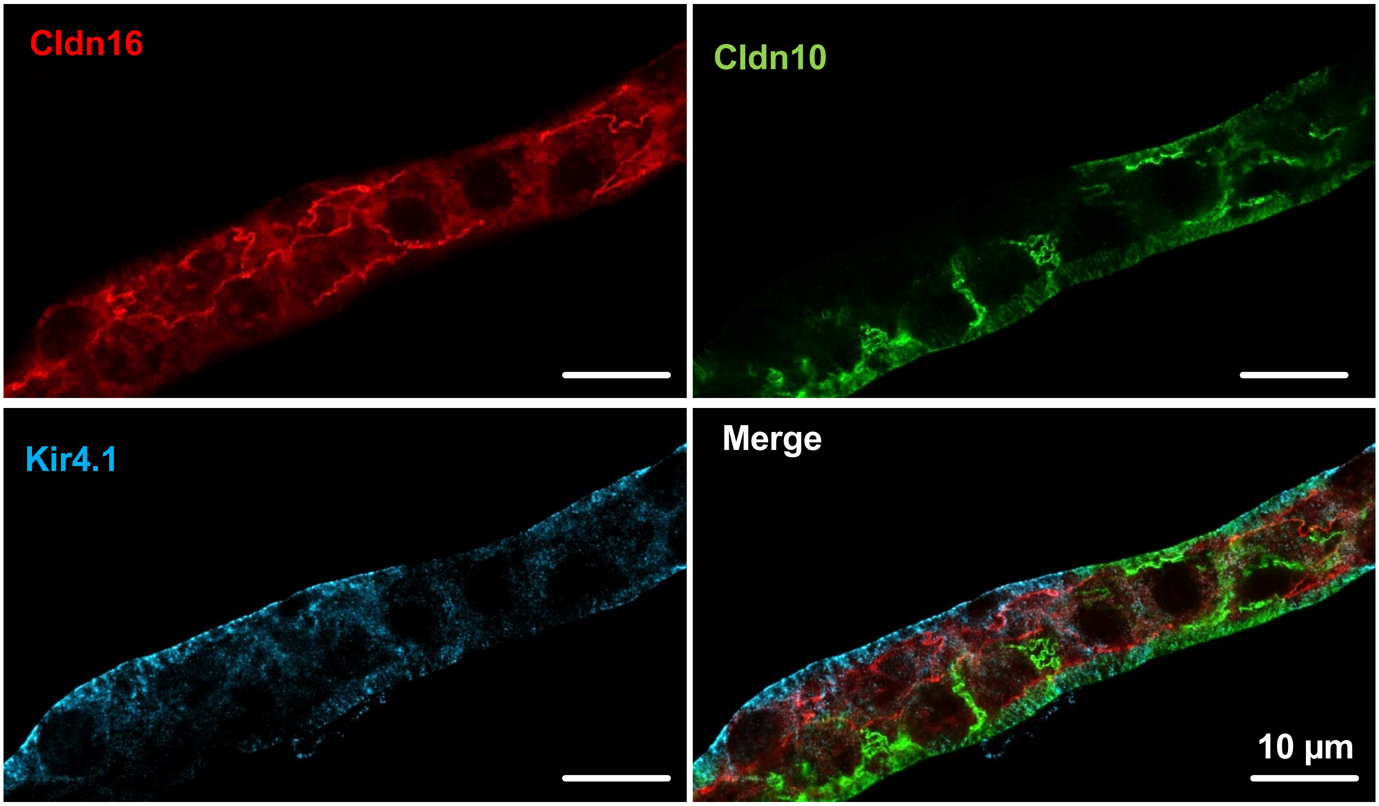

**Supplemental Figure 5. Cell type distribution and kidney zone localization of mouse TAL cells as determined by enriched single-nucleus RNA sequencing**

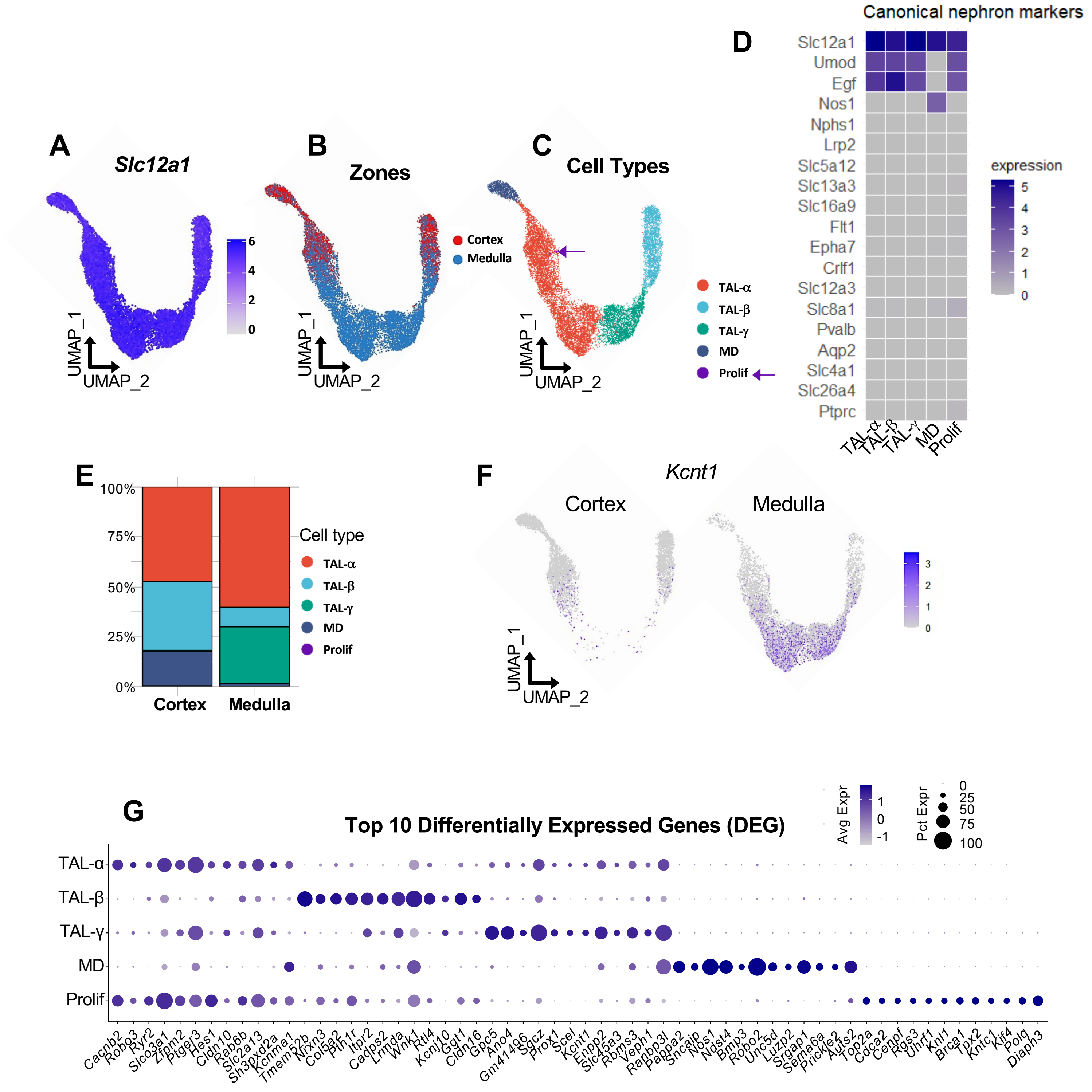

**Supplemental Figure 6. Highlighted gene expression of enriched single-Nnucleus RNA-Seq analysis from mouse**

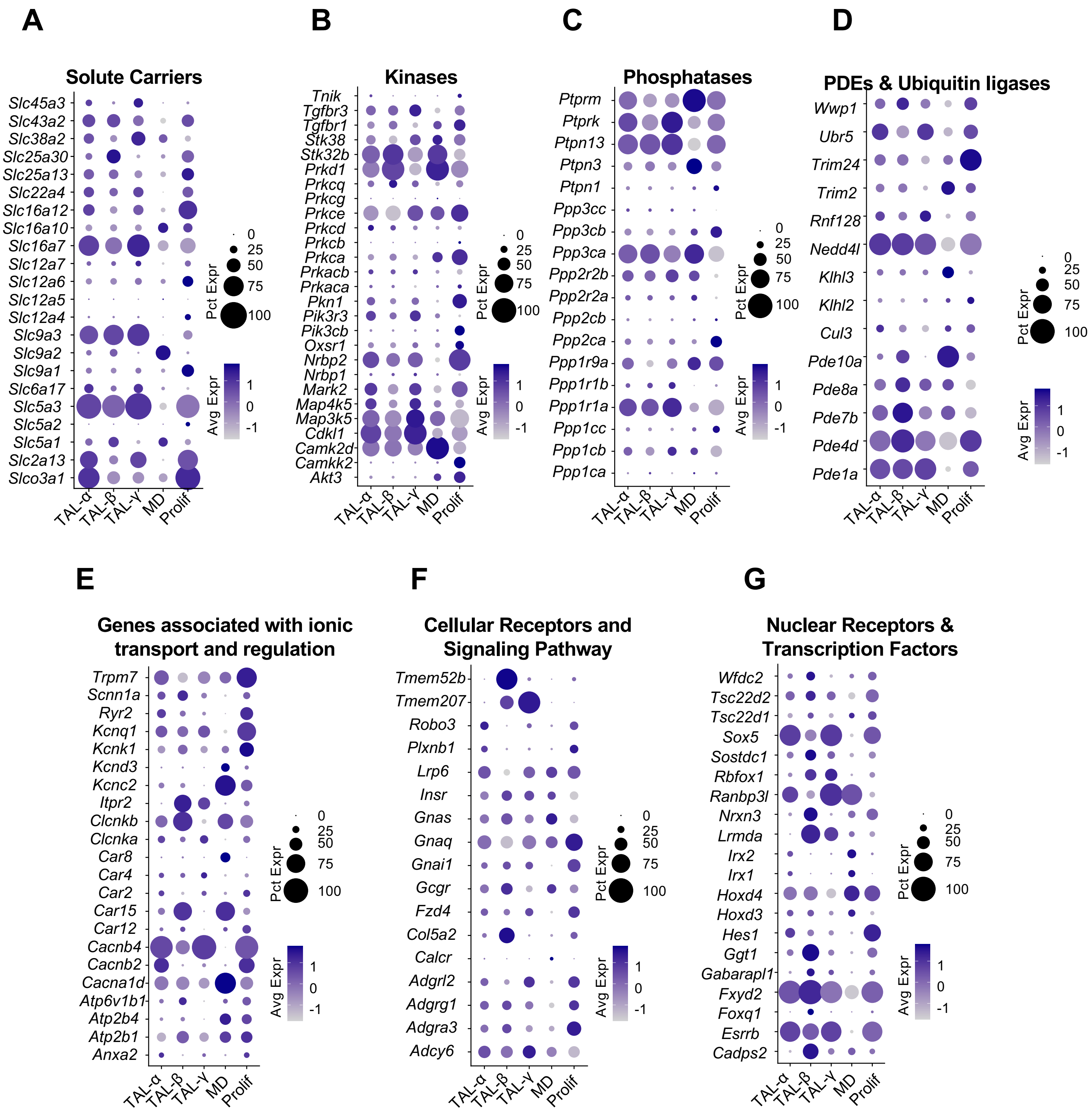

**Supplemental Figure 7. Cortico-medullary markers of enriched single-nucleus RNA-Seq analysis from mouse**

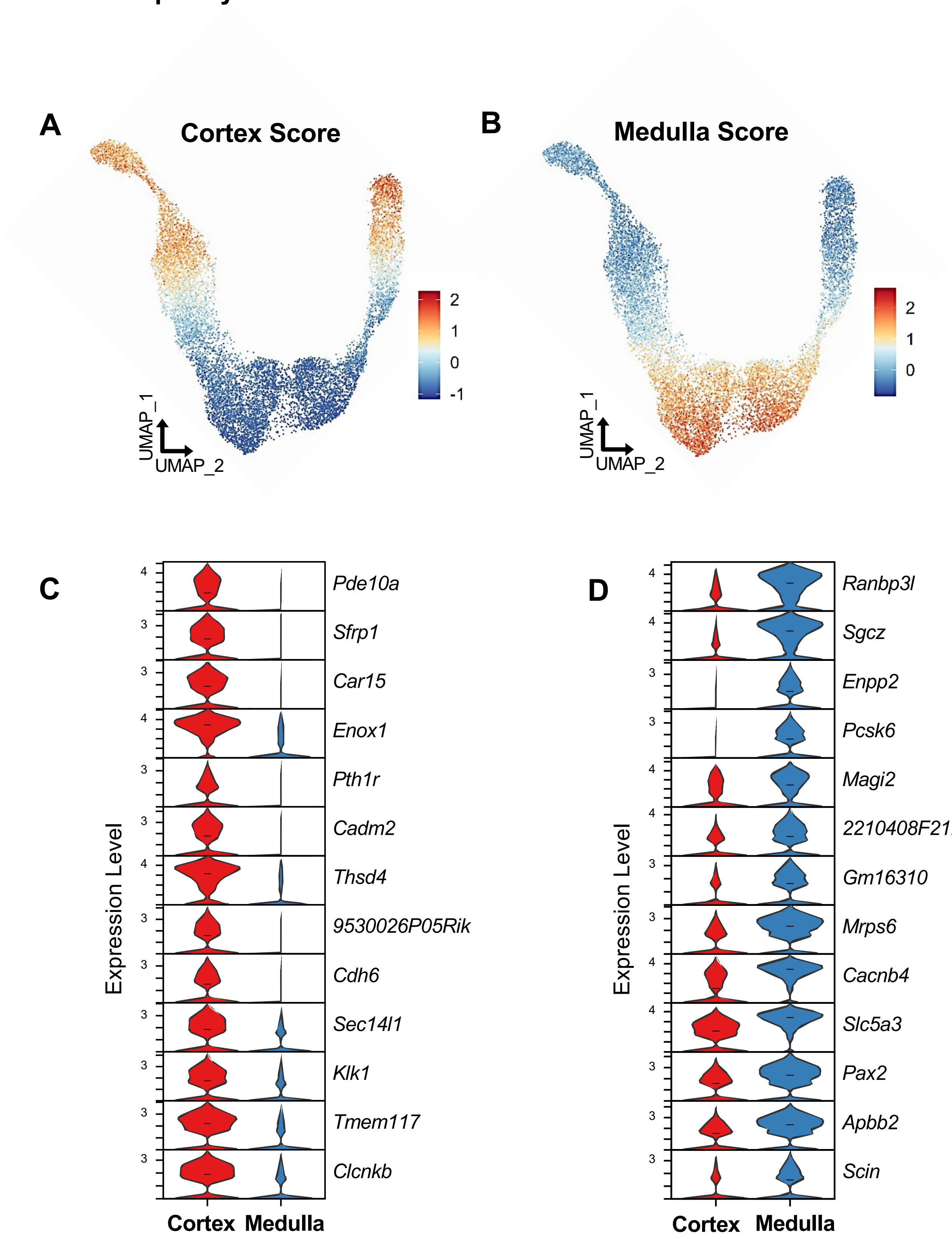

**Supplemental Figure 8. TAL cell diversity revealed by single-cell RNA-seq analysis, mouse whole-kidney previously published dataset**

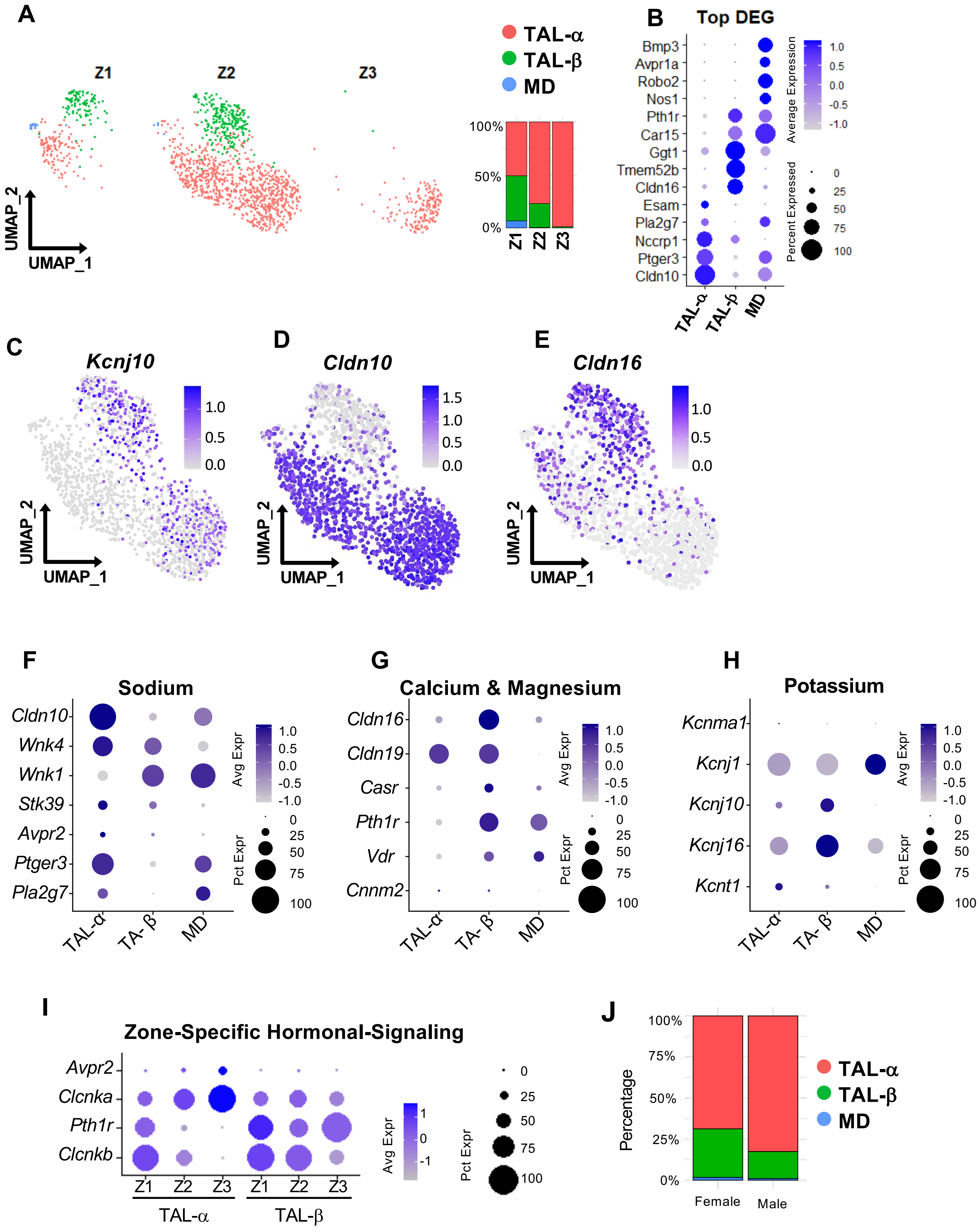

**Supplemental Figure 9. Human TAL cell population from KPMP data and comparison with mouse**

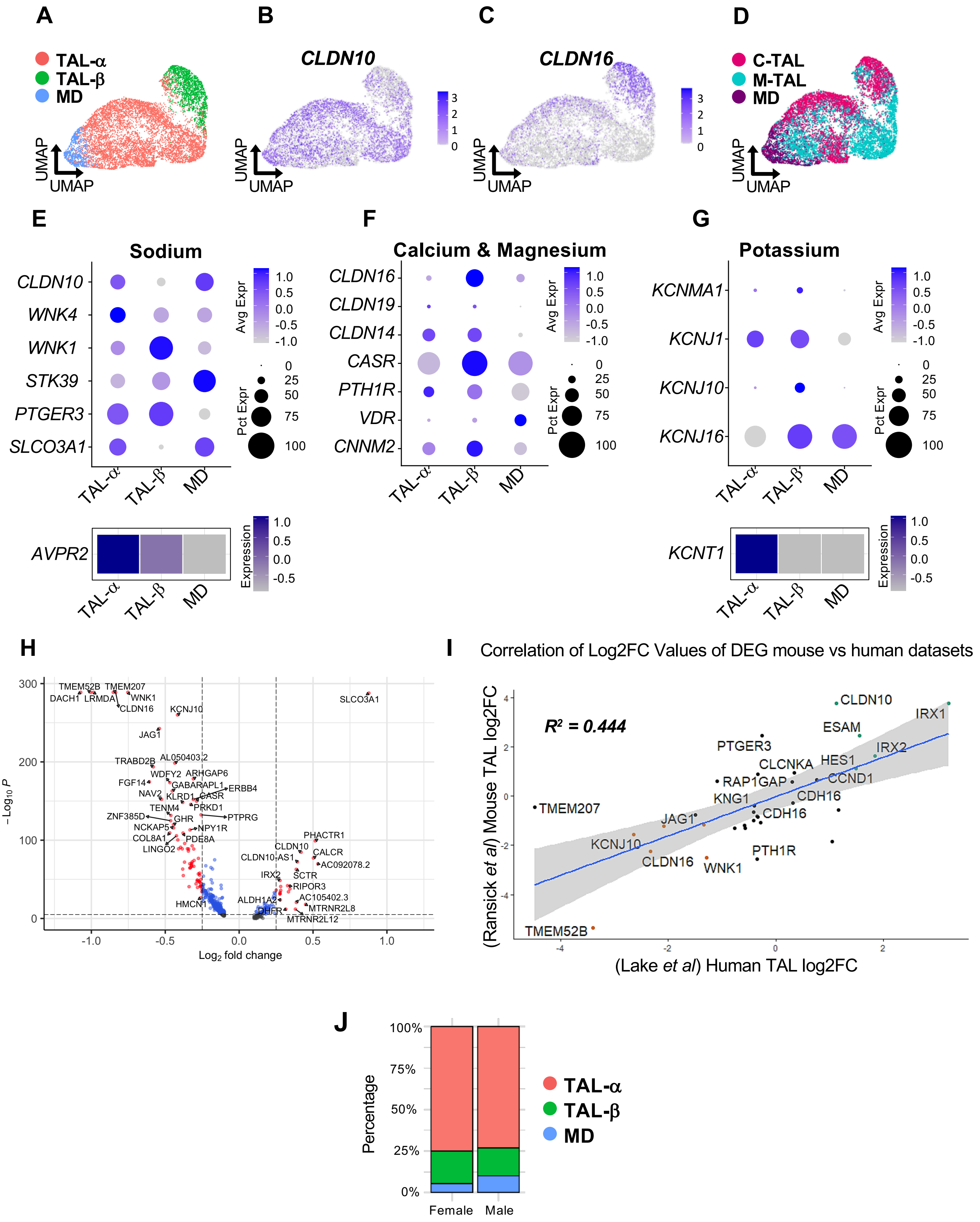

**Supplemental Figure 10. TAL cell diversity revealed by single-nuclei RNA-seq analysis, rat kidney and comparison with mouse**

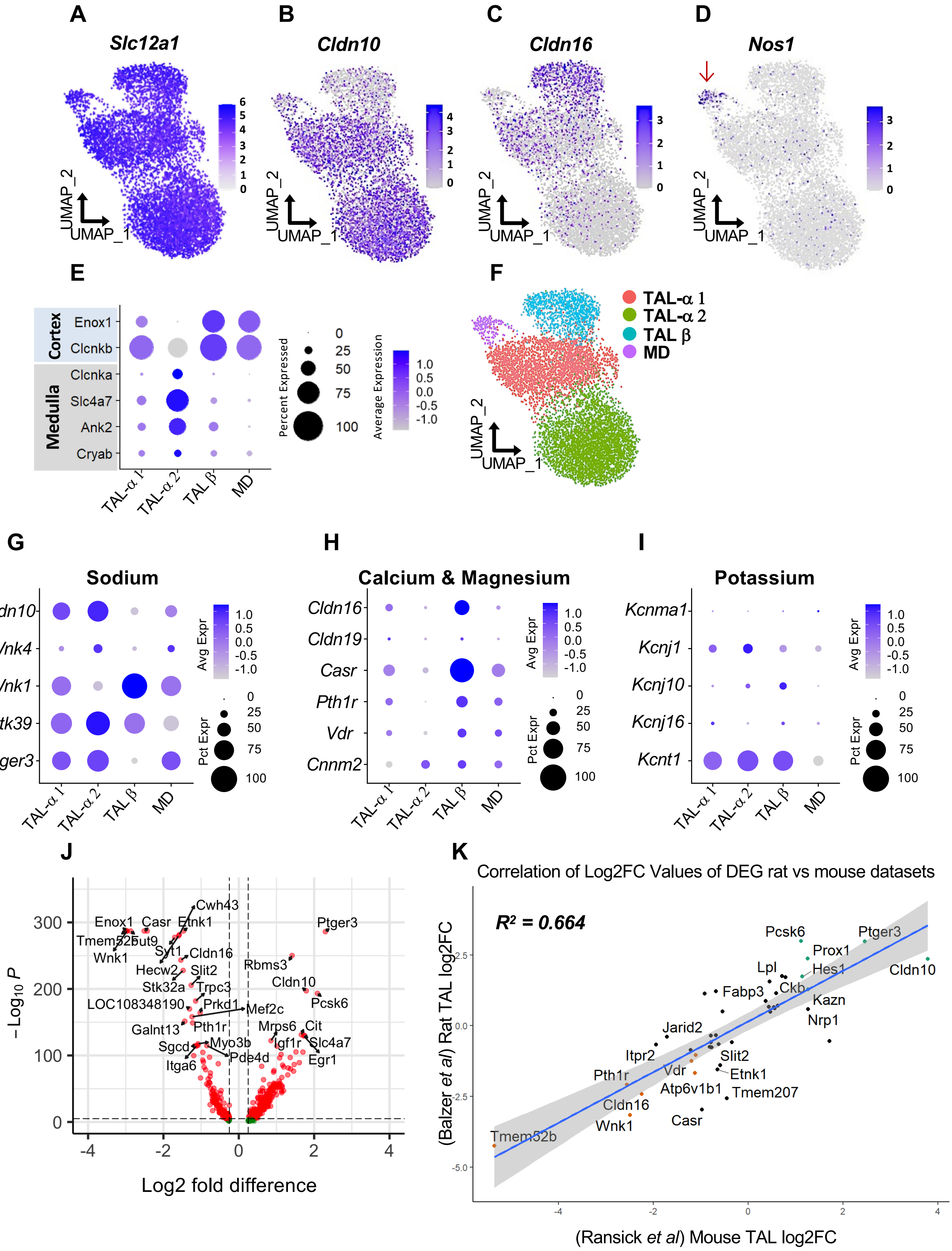

**Supplemental Figure 11. Anatomically stratified TAL cell type characterization from transcriptomic analysis and expression of key membrane proteins**

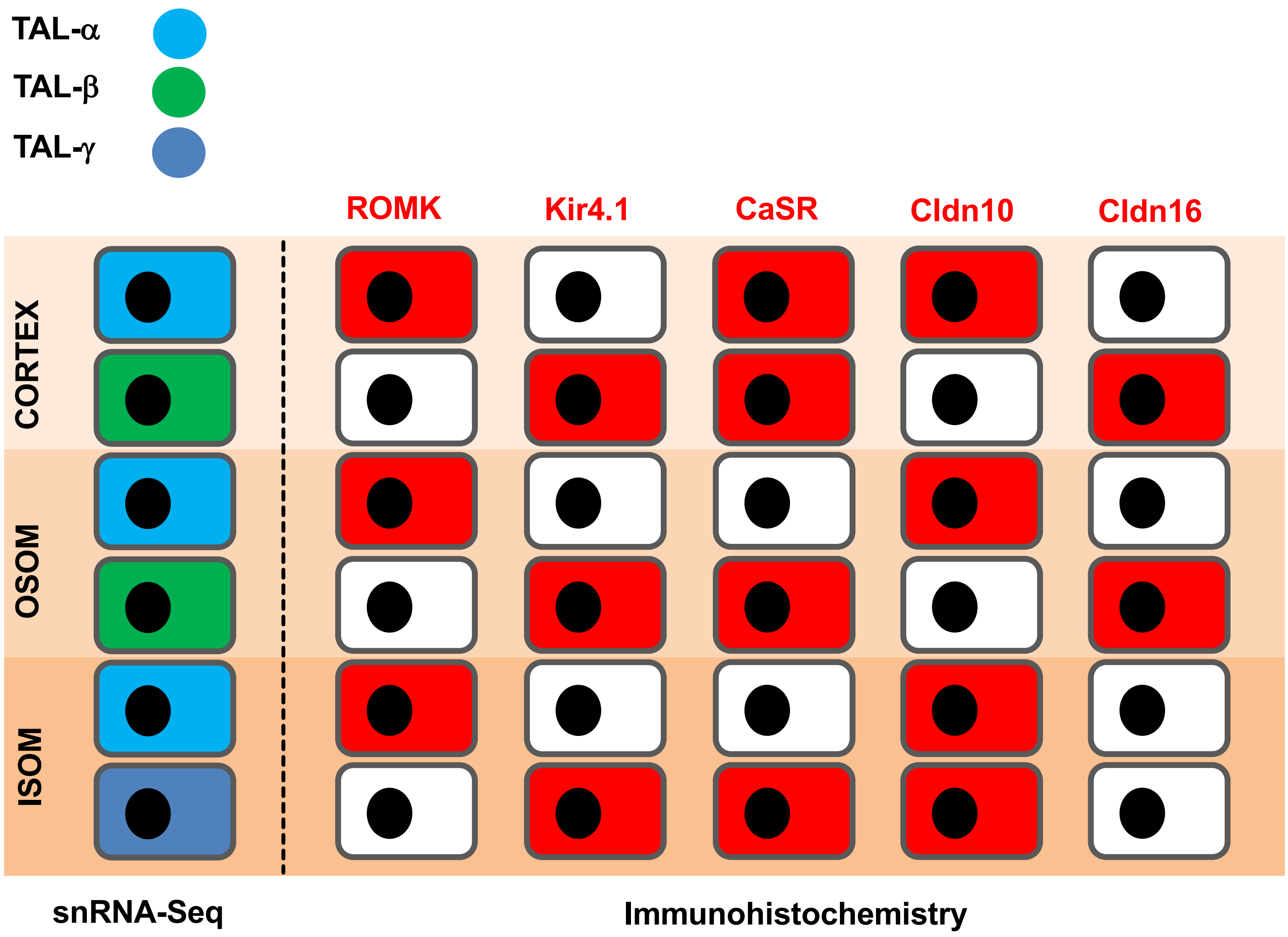


**Full Cell Type Classification**

| Name | ROMK | Kir4.1 | Claudin 10 | Claudin 16 | Region |
|---|---|---|---|---|---|
| TAL-α | + | - | + | - | All |
| TAL-β | - | + | - | + | Cortex & OSOM |
| TAL-γ | - | + | + | - | ISOM |

**Transcriptomic Cell Type Classification**

| Name | ROMK | Kir4.1 | Claudin 10 | Claudin 16 | Region |
|---|---|---|---|---|---|
| TAL-α | + | - | + | - | All |
| TAL-β | + | + | - | + | Cortex & OSOM |
| TAL-γ | + | + | + | - | ISOM |
